# Experimental Stroke Induces Chronic Gut Dysbiosis and Neuroinflammation in Male Mice

**DOI:** 10.1101/2020.04.29.069575

**Authors:** Allison L. Brichacek, Divine C. Nwafor, Stanley A. Benkovic, Sreeparna Chakraborty, Sophia M. Kenney, Maria E. Mace, Sujung Jun, Catheryne A. Gambill, Wei Wang, Heng Hu, Xuefang Ren, Jessica M. Povroznik, Elizabeth B. Engler-Chiurazzi, Donald A. Primerano, James Denvir, Ryan Percifield, Aniello Infante, Jennifer Franko, Rosana Schafer, Darren E. Gemoets, Candice M. Brown

**Affiliations:** Department of Microbiology, Immunology, and Cell Biology, School of Medicine; Department of Neuroscience, School of Medicine; Department of Physiology and Pharmacology, School of Medicine; Center for Basic and Translational Stroke Research, Blanchette Rockefeller Neuroscience Institute; Department of Biostatistics, School of Public Health; Genomics Core Facility, West Virginia University, Morgantown, WV 26506; Department of Biomedical Sciences, Marshall University Joan C. Edwards School of Medicine, Huntington, WV 25755

**Keywords:** stroke, microbiome, microbiota-gut-brain axis, neuroinflammation, gut dysbiosis, ischemia, stroke behavior

## Abstract

Recent literature implicates gut epithelia mucosa and intestinal microbiota as important players in post-stroke morbidity and mortality. As most studies have focused on the acute effects of stroke on gut dysbiosis, our study objective was to measure chronic, longitudinal changes in the gut microbiota and intestinal pathology following ischemic stroke. We hypothesized that mice with experimental ischemic stroke would exhibit chronic gut dysbiosis and intestinal pathology up to 36 days post-stroke compared to sham controls. Male C57BL/6J mice were subjected to 60 minutes of transient middle cerebral artery occlusion (tMCAO) or sham surgery. To determine the long-term effects of tMCAO on gut dysbiosis, fecal boli were collected pre- and post-tMCAO on days 0, 3, 14, and 28. Bioinformatics analysis demonstrate significant differences in abundance among Firmicutes and Bacteroidetes taxa at the phylum, family, and species levels in tMCAO compared to sham mice that persisted up to one month post-stroke. The most persistent changes in post-stroke microbial abundance were a decrease in bacteria family S24-7 and significant increases in *Ruminococcaceae*. Overall, these changes resulted in a persistently increased Firmicutes:Bacteroidetes ratio in stroke animals. Intestinal histopathology showed evidence of chronic intestinal inflammation that included marked increases in immune cell infiltration with mild-moderate epithelial hyperplasia and villous blunting. Increased astrocyte and microglial activity were also detected one-month post-stroke. These results demonstrate that acute, post-stroke disruption of the gut-brain-microbiota axis progresses to chronic gut dysbiosis, intestinal inflammation, and chronic neuroinflammation.

**Clinical Perspectives:**

- The microbiota-gut-brain axis, recently implicated in several neurological disorders, remains largely unexplored at chronic time points post-tMCAO.
- Our results demonstrate chronic gut dysbiosis, prolonged behavioral deficits, and persistent cerebral and intestinal inflammation post-tMCAO in male C57BL/6J mice.
- These results suggest that manipulation of microbiota may help reduce poor outcomes after stroke and lead to improved post-stroke functional recovery.

## Introduction

Ischemic stroke is an acute neurological disorder that induces both brain and systemic inflammatory immune responses. An emerging body of data from humans and mice strongly suggests that changes in gut microbiota are common during stroke and that these changes actively contribute to disease pathophysiology. Up to half of stroke patients experience gastrointestinal (GI) complications after injury, including gut dysbiosis, increased gut barrier permeability and hemorrhaging, dissemination of gut bacteria to other organs, and sepsis from gut-originating bacteria [1–3]. Consequently, stroke patients with these complications often experience a delayed stroke recovery time, higher rates of mortality, and worse neurological outcomes [4]. One critical barrier to progress in the stroke field is an incomplete understanding of the specific taxa that contribute to gut dysbiosis, as well as the mechanisms through which specific taxa affect the pathophysiology of stroke within the microbiota-gut-brain axis.

The microbiota-gut-brain axis is a bidirectional communication system through which gut microbes mediate communication between the brain and the gut [5]. The gut microbiota modulates normal gut physiology and signaling along this axis to promote homeostasis [6,7]. Communication between the gut microbiota and the central nervous system (CNS) integrates neural, immune, endocrine, and metabolic pathways [8,9]. Gut microbiota can modulate the release of neurotransmitters by regulating the levels of their precursors, or they can synthesize those products themselves. Short-chain fatty acids (SCFAs), the metabolic products of microbial species, mediate crosstalk along this axis directly via humoral immunity and indirectly via the autonomic nervous system, hormonal routes, and immune signaling pathways [5,10]. Loss of homeostasis of the gut microbiota is linked to obesity, metabolic disorders, and neurological dysfunction. This can culminate into behavioral changes, including those associated with depression and anxiety, which are associated with stroke [11,12].

The majority of clinical and animal stroke microbiome studies have focused on early time-points, typically spanning ≤7 days [13–15]. Clinical studies exploring changes in the gut microbiome after stroke have shown significant dysbiosis in stroke patients. The stroke microbiota population includes an increased abundance of opportunistic pathogens and fewer commensal bacteria [16]. Changes in the microbiome may also be a risk factor leading to disease, including stroke and stroke-related comorbidities. A prospective study showed that opportunistic pathogens, including *Enterobacteriaceae* and *Veillonellaceae*, and lactate-producing bacteria, including *Lactobacillus* and *Bifidobacterium*, were increased in patients who had a high risk of stroke compared to low risk controls [17]. Changes in the microbiome of at-risk or post-stroke patients can also be mimicked in rodent models. Several groups have observed changes in species diversity and taxa abundance in mice post-injury [13,14].These short-term changes in the gut microbiota after stroke are coupled to increased intestinal permeability, T-cell proliferation, and harmful IL-17-producing γδ T-cells that migrate to the brain [6,13,18]. Spontaneously hypertensive stroke prone rats (SHRSP) showed increased concentrations of *Lactobacillus* and an increased Firmicutes:Bacteroidetes ratio compared to normotensive controls [19]. In contrast, much less is known about the long-term effect(s) that stroke has on microbial changes in the gut and how these changes may contribute to the severity of stroke-associated neuropathology and functional outcomes.

The goal of this study was to test whether there are longitudinal, sustained alterations in the gut microbiome following ischemic stroke, and how these changes are associated with long-term changes in innate immunity, neuroinflammation, and behavior. In this study, we hypothesized that there would be a persistent, long-term loss of diversity in the gut microbiome following ischemic stroke, and that the decreased microbial diversity would be paralleled by increased brain and systemic inflammation. To address this, we employed a transient middle cerebral artery occlusion (tMCAO) model in C57BL/6J male mice to explore longitudinal changes in the microbiome at baseline (pre-stroke), 3, 14, and 28 days post-stroke compared to sham-injured controls. Our results demonstrate that there is a long-term increase in the Firmicutes:Bacteroidetes ratio that is coupled with the loss of several beneficial bacteria taxa after stroke. These changes were accompanied by sustained neuroinflammation and intestinal pathological changes associated with gut dysbiosis. A greater understanding of which harmful bacteria persist and how these can be targeted may lead to improved therapeutic options for individuals who experience an ischemic stroke.

## Materials and Methods

### Animals

Male C57BL/6J, i.e. wild type (WT), mice were obtained from The Jackson Laboratory (Bar Harbor, ME, USA). All animals were bred and housed in the West Virginia University Office of Laboratory Animal Resources vivarium facilities. Animals used in experiments were 3-6 months of age and were randomly allocated to experimental groups. Mice were group-housed according to experimental group (sham vs. stroke) in environmentally controlled conditions with a reverse light cycle (12:12 hr light/dark cycle) and provided food and water *ad libitum*. Whenever possible, the experimental design was modeled on ARRIVE [20] and STAIR [21] guidelines. All experiments were approved by the Institutional Animal Care and Use Committee of West Virginia University.

### Transient Middle Cerebral Artery Occlusion

Transient middle cerebral artery occlusion (tMCAO) was carried out as previously described [22]. Briefly, surgical anesthesia was induced using 4-5% isoflurane, then maintained at 1-2% via face mask. Rectal body temperature was maintained at 37±0.5°C during the surgical procedure. Focal cerebral ischemia was performed via a 60-minute occlusion of the right middle cerebral artery (tMCAO) with a 6.0 monofilament suture (Doccol, Sharon, MA). Measurements of cerebral blood flow (CBF) were taken at baseline, immediately following insertion of the monofilament (ischemia), and immediately following removal of the monofilament (reperfusion). Briefly, the skull was exposed with a small incision and CBF images were acquired using a MoorFLPI laser Doppler speckle flowmetry system (Moor Instruments, England). Flux values were measured at the three time points (prior to ischemia, i.e. baseline; immediately post-ischemia; and immediately post-reperfusion) by comparison of a 3.8 cm × 3 cm area of the ipsilateral (right) and contralateral (left) hemispheres. Fold change was calculated by normalizing ischemic and reperfusion values to baseline. To prevent hypothermia, animals were placed in an enclosed room maintained at a temperature of 73°C for the first 24 hr post-tMCAO. All animal weights and temperatures were monitored and recorded for the first three days following sham or tMCAO surgery, and all animals received fresh moist chow supplemented with Nutri-Cal (Vetoquinol, Fort Worth, TX) daily for four days post-surgery. For analgesia, all mice received a daily bupivacaine injection (2.5 mg/ml) (Hospira, Inc., Lake Forest, IL) subcutaneously near the neck wound for the first three days after surgery.

### Experimental Design for each tMCAO Cohort

#### Cohort 1 - 24-hour Cohort

Baseline weights were collected prior to tMCAO or sham surgery (tMCAO n=6, sham n=6 total from 3 independent studies). Animals in this cohort were euthanized approximately 24 hrs post-surgery. Serum was collected via cardiac puncture. Brain tissue was collected following saline perfusion for flow cytometric analysis.

#### Cohort 2 - 31-36-day Cohort

Baseline weight and fecal boli samples were collected prior to tMCAO or sham surgery on experimental day 1 (tMCAO n=7, sham n=7 total from 3 independent studies). Modified neurological severity scores, weights, and temperatures were assessed on days 2, 3, 7, 14, 21, and 28. Behavioral testing was performed on days 3 and 7, then once weekly for four weeks total. Additionally, fecal boli were collected on days 3, 14, and 21. Animals in this cohort were euthanized between 31 and 36 days post-surgery, and serum and tissues were collected. The experimental design is summarized in **Figure 1**.

**Fig. 1.**
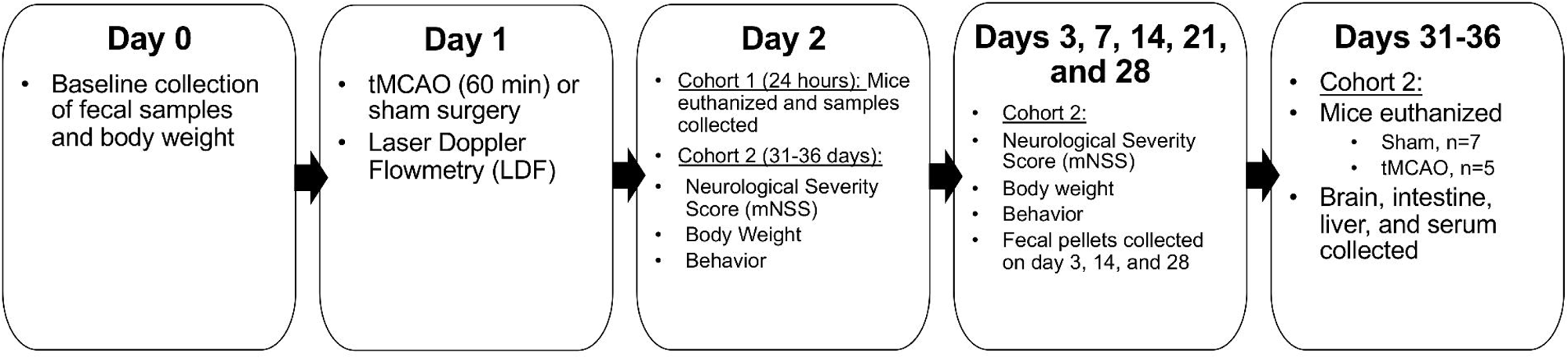
Experimental schematic for mice in two cohort studies. Prior to the start of the study, baseline weights and fecal boli samples were taken for longitudinal comparison. Male C57BL/6J mice were randomly assigned to sham or stroke (tMCAO; 60 min) surgery, confirmed using LDF. Cohort 1 animals (sham, n=6; stroke, n=5) were euthanized 24 hours after surgery, and tissues were collected for analysis. Cohort 2 animals were part of a longitudinal study where fecal boli, behavior, and clinical scores were taken at least once weekly. Cohort 2 mice (sham, n=7; stroke n=5) were euthanized between days 31-36 post-surgery (∼35 days) followed by collection of serum, brain, intestine, and liver for analysis.

### Clinical Scores and Behavioral Testing

Clinical severity following surgery was assessed on experimental days 2, 3, 7, 14, 21, and 28, using a murine modified neurological severity score (mNSS) adapted from Chen *et al.* [23]. Briefly, mice underwent a four-part test which examined (1) reflex, (2) walking ability, (3) balance, and (4) head and limb flexion. Individual test scores were summed for a total score of 0 (normal/unimpaired) to 14 (severely impaired). Behavioral testing was performed on experimental days 3, 7, 14, 21, and 28. Open-field testing was used to assess locomotor activity. Mice were placed into 16 × 16 × 15 inch chambers (Photobeam Activity System, San Diego Instruments, San Diego, CA) for a total of 20 min. Activity was measured every 2.5 mins by recording the number and location of beam breaks using a 3 × 3 inch center. The number of horizontal (x-, y-direction), central, and rearing (z-direction) movements were recorded. Rotarod testing was used to assess locomotor function. Mice were subjected to accelerating speeds of 4 to 40 rpm for a maximum of 300 sec; each mouse underwent a total of three rotarod trials per testing time point.

### Sample Collection and Processing

Mice were deeply anesthetized with isoflurane and blood was collected via cardiac puncture into Microvette tubes (Sarstedt; Nümbrecht, Germany). Serum was isolated via centrifugation at 10,000 x g for 5 mins. Animal tissues were then perfused transcardially with 10 ml of 0.9% saline with a perfusion pump (Masterflex 7524-10, Cole-Parmer, Vernon Hills, IL) set to 5.0 ml/min. Livers were harvested and stored in RNA later at −20°C. The remaining tissues were fixed with 50 ml of 4% cold paraformaldehyde (PFA) (Fisher Scientific, Pittsburgh, PA) and stored at 4°C in 4% PFA. Brain and intestine (duodenum) tissues were rinsed in 0.01 M phosphate buffered saline (1X PBS) and incubated sequentially in 10%, 20%, and 30% sucrose in 1X PBS for 24 hr each.

### Flow Cytometry

Single cell suspensions were obtained as previously described [24]. Briefly, brains were perfused and collected as described above. Whole brains were removed from the skull, separated into ipsilateral and contralateral hemispheres, separately minced, and homogenized mechanically into respective Gentle MACS C tubes containing 50 µL enzyme P + 1900 µL buffer Z and 20 µL of Buffer Y + 10 µL of Enzyme A (Miltenyi Biotec, Auburn, CA). Samples were passed through a 70 µm cell strainer (Millipore, Temecula, CA) to obtain single cell suspensions. Cells were then collected by centrifugation at 300 g for 10 min at 4°C. Red blood cells were lysed with 4 ml of ACK Lysis Buffer (Lonza, Walkersville, MD) per brain, incubated for 2-3 min at room temperature and washed. Optionally, cells were passed through an additional 70 μm filter. Once in a single cell suspension, CD45+ immune cells were isolated using magnetic labeling and the autoMACS (Miltenyi Biotec, Auburn, CA). CD45-cells were stored at 4°C, while CD45+ cells were stained for flow cytometry. Viability and total brain cell yield was determined by trypan blue exclusion and re-suspended to a final concentration of 2.5-3 × 10^6^ cells/ml with FACS buffer. Cells were washed twice in sodium azide and protein-free cold 1X PBS and stained with fixable viability dye eFluor780 (eBioscience, San Diego, USA) for 30 min at 4°C in dark. Cells were briefly washed with FACS buffer containing 5 mM EDTA, 2% FBS in cold PBS, and blocked with Ultra-Leaf purified anti-mouse CD16/32 (Clone: 93; BioLegend, San Diego, CA) for at least 20 min. Following non-specific blocking, cells were stained with monoclonal antibodies for CD45-PE (Clone: REA737), CD11b-VioBlu, CD11c-PerCp-Vio700, Ly6G-APC, Ly6C-PE-Vio770 (Miltenyi Biotec, Auburn, CA) or anti CD4-FITC (Clone: RM4-5), CD25-PE (Clone: PC61.5), CD19-PerCp-Cy5.5 (Clone: 1D3) and CD8a-eFluor450 (Clone: 53-6.7; eBioscience, San Diego, USA) for 10 min at 4°C. Appropriate single stained controls were prepared for fluorophore compensation. For intracellular FOXP3 detection, following staining for the surface markers (CD4 or CD25), cells were permeabilized using the mouse T regulatory cell staining kit (Invitrogen, USA) and stained with anti-FOXP3-PE-Cy5 (Clone: FJK-16s) according to the manufacturer’s instructions. Cellular fluorescence was quantified with a BD LSR Fortessa using FACS Diva software (BD Biosciences, San Jose, CA). Single cells were identified by forward scatter and side scatter, and viable cells were gated. Cells were gated for CD45^+^ populations and then divided into lymphoid cells, which included B cells (CD11c^-^, CD19^+^), T-regulatory cells (CD4^+^, CD25^+^, and Foxp3^+^); myeloid cells, which included neutrophils (Ly6G^+^), monocytes (Ly6C^+^), infiltrating macrophages (CD45^hi^, CD11c^-^, and CD11b^+^); and brain residential microglia (CD45^lo^, CD11b^+^). All data were compensated and spectral overlap was minimized using the automatic compensation method of BD FACS Diva software (BD Biosciences, San Jose, CA).

### Cytokine Array

Serum (100 μl) collected from mice in cohorts 1 and 2 was shipped on dry ice to RayBiotech, Inc. (Norcross, GA) for Quantibody testing using a custom 8-cytokine array to determine the concentration of the following cytokines (limits of detection in parentheses in pg/ml): monocyte chemoattractant protein 1 (MCP-1 or CCL2, 5.5 – 4,000), P-selectin (12.6 – 4,000), tumor-necrosis factor alpha (TNF-α, 4.0 – 10,000), vascular endothelial growth factor (VEGF, 8.3 – 4,000), endothelial-leukocyte adhesion molecule 1 (E-selectin, 14.3 – 4,000), L-selectin (38.8 – 10,000), vascular cell adhesion protein 1 (VCAM1, 8.2 – 4,000), and intracellular cell adhesion protein 1 (ICAM1, 19.5 – 10,000). Sample concentrations were calculated based on a standard curve and reported in pg/ml.

### Measurement of serum nitric oxide

Serum NOx (nitrite + nitrate) are stable intermediates of nitric oxide (NO) in biological solutions. Total NOx was measured by ozone-based chemiluminescence using a Sievers 280i Nitric Oxide Analyzer (Boulder, CO) as previously described [25,26]. Data were acquired using Liquid Program software v 3.22 PNN, Sievers. All standards and diluted samples were run in duplicate; sample concentrations (μM) were interpolated from the standard curve for all chemiluminescence values and the mean was reported.

### RNA Isolation and quantitative real time PCR

Liver tissues were harvested as described above and saved in RNA*later* (Invitrogen, Carlsbad, CA) at −20 °C. Tissues were homogenized and total RNA was isolated using RNeasy Mini Kit (Qiagen, Germantown, MD) standard protocol. RNA concentrations were quantified using a Nanodrop spectrophotometer (ThermoFisher Scientific, Carlsbad, CA). 2.5 μg template RNA was used to make cDNA using SuperScript IV VILO Master Mix (ThermoFisher Scientific, Carlsbad, CA). PCR was completed using a custom TaqMan Well FAST Plate (Applied Biosystems, Foster City, CA) array of 48 different mouse genes using a Step One Plus RT PCR system (Applied Biosystems, Foster City, CA) modified based on a previously published array of oxidant/antioxidant genes; *Hprt* (hypoxanthine transferase), *Gusb* (glucuronidase beta), *and Gapdh* (glyceraldehye-3-phosphate dehydrogenase) were employed as housekeeping genes [27]. Results were analyzed using DataAssist 3.01 (Applied Biosystems).

### Sectioning and histology for brain and intestine tissues

Brains and intestines were mounted onto the freezing stage of the sliding microtome (Model HM450, ThermoScientific, Waltham, MA). Brain and intestine sections were cut at 35μ, collected into PBS+azide (0.6g sodium azide/liter PBS), and stored at 4°C until use. *Periodic Acid Schiff (PAS) Staining*: Intestine sections were mounted onto microscope slides and allowed to air dry overnight. The sections were stained for PAS using a commercial kit (ES3400-IFU, Azer Scientific, Morgantown, PA) with the following modifications: no deparaffinization; room temperature water used for washes; hematoxylin counterstain for one minute; no running tap water, use 2 – 3 changes in staining dish.

### Immunohistochemistry

Mouse brains were analyzed as free-floating sections to localize immunoreactivity for GFAP (Z0334, Dako/Agilent; 1:10,000 primary, 1:10,000 secondary), Iba-1 (019-19741, Wako; 1:2,000 primary, 1:1,000 secondary), ALPL (702454, Invitrogen, 1:100 primary, 1:1,000 secondary), and CD11b (ab 133357 Abcam, 1:500 primary, 1:1,000 secondary) using a modified ABC procedure (Vector Laboratories, Burlingame, CA) as previously described [28,29]. Briefly, sections were treated to remove endogenous peroxidases followed by permeabilization. Sections were transferred into the appropriate primary antibody solution and incubated overnight. Sections were then incubated in secondary antibody for two hours the following day. Sections were then incubated with Avidin D-HRP (for GFAP, 1:1000 in PBS, Vector Laboratories, Burlingame, CA) or ABC reagent (for Iba-1 and CD11b) for 1h at room temperature. Next, antigens were visualized with 3-3’ diaminobenzidine (DAB) as the chromogen. Lastly, sections mounted onto microscope slides, (Unifrost+, Azer Scientific, Morgantown, PA), air dried overnight, dehydrated through a standard ethanol series, and cover slipped with Permount (Fisher Scientific, Pittsburgh, PA)

### Microscopy and image analysis

Slides were viewed on a Leica DM6B microscope and images were captured using Leica LASX software (Leica Microsystems, Buffalo Grove, IL). For GFAP, images were captured from cortex of three adjacent sections at 40X. A batch processing file was programmed in Photoshop with the following steps: (1) convert to grayscale, (2) invert, (3) threshold = 144, (4) select all, (5) record measurement – mean gray scale. Mean intensity values were collected and analyzed by ANOVA using Prism 8 (GraphPad Software, Inc., La Jolla, CA). Iba-1 images were captured from medial orbital cortex of three adjacent sections at 20X. A batch processing file similar to that for GFAP was programmed in ImageJ version 2.0. Cell bodies and cell area were quantified manually using the count tool and were recorded and analyzed in Prism 8 (GraphPad Software, Inc., La Jolla, CA).

### 16S rRNA Sequencing

Fecal pellets were collected on experimental days 0 (day before surgery), 3, 14, and 28. DNA was extracted using the DNeasy PowerSoil Kit (Qiagen Inc., Valencia, CA). Samples (n=5-7/group/time point) were processed using a standard protocol with minor modifications. Instead of vortexing, samples were homogenized (Mini-Beadbeater, Biospec Products; Bartlesville, OK) at maximum speed for 5 mins. Sample DNA concentrations were measured on a Qubit fluorometer (ThermoFisher Scientific; Carlsbad, CA), followed by PCR amplification of the V3-V4 hypervariable region of the bacterial 16S rRNA gene. Library preparation of samples was performed using a two-step dual-indexing strategy modified from Fadrosh *et al.* [30]. Amplicon size and overall quality were assessed on High Sensitivity Bioanalyzer Chips (Agilent, Santa Clara, CA). Following library preparation, amplicons were sequenced in the Rapid Run mode on an Illumina HiSeq1500 instrument in a 2×270 paired end run with standard Illumina sequencing protocols. A total of 236 samples were included in the run, but only 107 samples were included as part of this study. Illumina run metrics indicated that 271,905 “contigs” (= 543,810 individual reads per sample) passed quality control filters.

### Bioinformatics and Biostatistics

A custom script was used to remove heterogeneity spacers within primers as described in Fadrosh *et al.* [30]. Within the Dada2 framework, reads were quality filtered to an expected error rate of 2, as well as length trimmed [31]. After learning error rates and doing sample inference, the resulting sequences were filtered by length and chimeras removed. Taxonomy was assigned using GreenGenes at 97% [32]. Taxa with differential abundance were identified using DESeq2, and population differences were explored using the vegan R package [33]. Further analyses were performed using alpha diversity, beta diversity, and abundance measures as response variables. For alpha diversity, analysis of variance/covariance was employed with Shannon diversity measures as the response. For beta diversity non-metric multidimensional (NMDS) scaling for ordination was employed via the vegan R package with the Bray-Curtis dissimilarity/distance metric. Significance tests involving beta diversity were performed using permutation ANOVA (PERMANOVA) in the vegan package. Relative abundances were found using amplicon sequence variants (ASVs), which were then converted to simple proportions for each operational taxonomic unit (OTU) rather than rarefaction [34] within each sample. In plots with abundance aggregated over a treatment the average proportion was calculated across samples to ensure that each sample was equally weighted in the aggregate. Abundance measures for longitudinal data sets were analyzed using a linear mixed-effects model with arcsine square root transformation (LMER); this statistical method can accommodate missing observations due to dropout, i.e. mice that died before the completion of the study [35]. Longitudinal data sets were also measured using a two-part zero-inflated regression model with random effects (ZIBR); this model cannot accommodate missing observations [36]. Adjusted p-values were reported for all results from both statistical models, and all analyses were performed using R statistical software [37], assuming a 0.05 significance level.

### Statistics

Data were analyzed by parametric or non-parametric (for skewed datasets) statistics as appropriate. Student’s t-test was used for pairwise comparisons in datasets with one independent variable, and one-way analysis of variance (ANOVA) was used to compare 3 or more groups in datasets with one independent variable. Two-way ANOVA was used to compare main effects or interactions between two independent variables. Post-hoc testing was used to compare differences between groups following ANOVA. Statistical details for individual experiments are described in the figure legends. Statistical analyses were performed in GraphPad Prism 8.0 or R (3.5.1) with p<0.05 as significant; all results are expressed as means ± SEM.

## Results

### Cerebral blood flow is suppressed post-tMCAO

Our experimental design included an examination of immune changes at early (24 hr, cohort 1) and late (31-36 days, cohort 2) time-points after stroke **(Figure 1)**. All animals received either transient middle cerebral artery occlusion (tMCAO; 60 mins) or sham surgery, validated by laser speckle flowmetry (**Figure 2**). A representative image of cerebral blood flow (CBF) at baseline (before tMCAO), after ischemia, and during reperfusion are shown (**Figure 2a-c**). Flux fold change comparisons between the ipsilateral and contralateral hemispheres at baseline, ischemia, and reperfusion showed that CBF was decreased in both hemispheres following tMCAO with a larger decrease, as expected, in the ipsilateral hemisphere (**Figure 2d**). The recovery of CBF following reperfusion was also smaller in the ipsilateral hemisphere compared to the contralateral hemisphere. These results confirm that our tMCAO model performed as expected during the ischemia and reperfusion phases of injury.

**Fig. 2.**
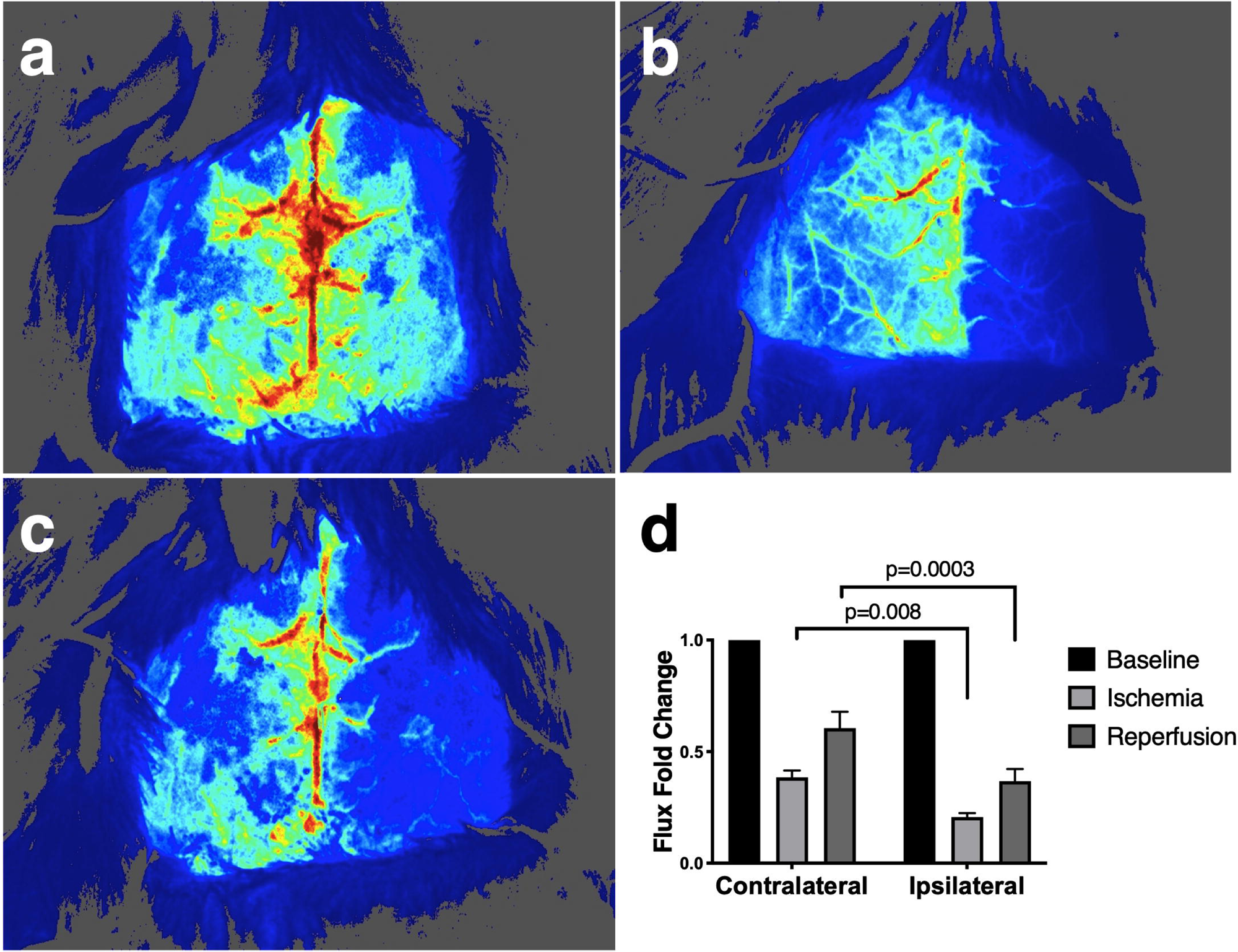
Cerebral blood flow (CBF) is decreased following 60 minutes of transient middle cerebral artery occlusion. Transient middle cerebral artery occlusion (tMCAO) was preformed via insertion of a microfilament for 60 mins. Representative laser speckle flowmetry images show cerebral blood flow for mice at baseline (**a**, before tMCAO), ischemia (**b**), and reperfusion **(c)**, where the ipsilateral hemisphere is on the right and the contralateral hemisphere is on the right. CBF for the ischemia and reperfusion phases were normalized to baseline and reported as flux fold change. The flux fold change between mice show a reduction in blood flow during ischemia and reperfusion in both hemispheres compared to baseline (main effects of hemisphere, time p<0.0001; hemisphere x time interaction p=0.01) with lower CBF in ipsilateral hemisphere compared to the contralateral hemisphere during ischemia (p=0.008) and reperfusion (p=0.0003) **(d)**. Flux fold change was analyzed by two-way ANOVA (main affects) with Sidak’s multiple comparison’s test (p-values shown on graphs) and reported as means ± SEM; n=13 tMCAO mice combined from both cohorts.

### Brain immune cell populations are elevated in the ipsilateral hemisphere post-tMCAO

Leukocyte infiltration is a critical immune event in early stroke. Whole brains were harvested at 24 hours post-MCAO or sham surgery. Brains were divided into ipsilateral and contralateral hemispheres for subsequent analysis. Following isolation of CD45+ cells, flow cytometric analysis was used to identify specific immune cell populations. Cell type counts from the ipsilateral hemisphere were normalized to the contralateral hemisphere, i.e. I:C ratio, to account for individual differences among mice. Results demonstrated a trending increase in the I:C ratios for stroke-injured mice compared to sham controls for CD11b+ monocytes/macrophages and CD3+ T cells (**Figure 3a-b**). In contrast, analysis of CD4+ and CD8+ T cells demonstrate an increase at 24 hours after tMCAO (**Figure 3c-d**). Additional analyses also determined that Ly6C+ monocytes, Ly6G+ neutrophils, and CD19+ B cells were not significantly different between sham and tMCAO groups (**Supplemental Figure 1**). The post-stroke infiltration of innate and adaptive immune cells into the ipsilateral hemisphere demonstrates that our model exhibits a prototypical proinflammatory response at 24 hours post-stroke.

**Fig. 3.**
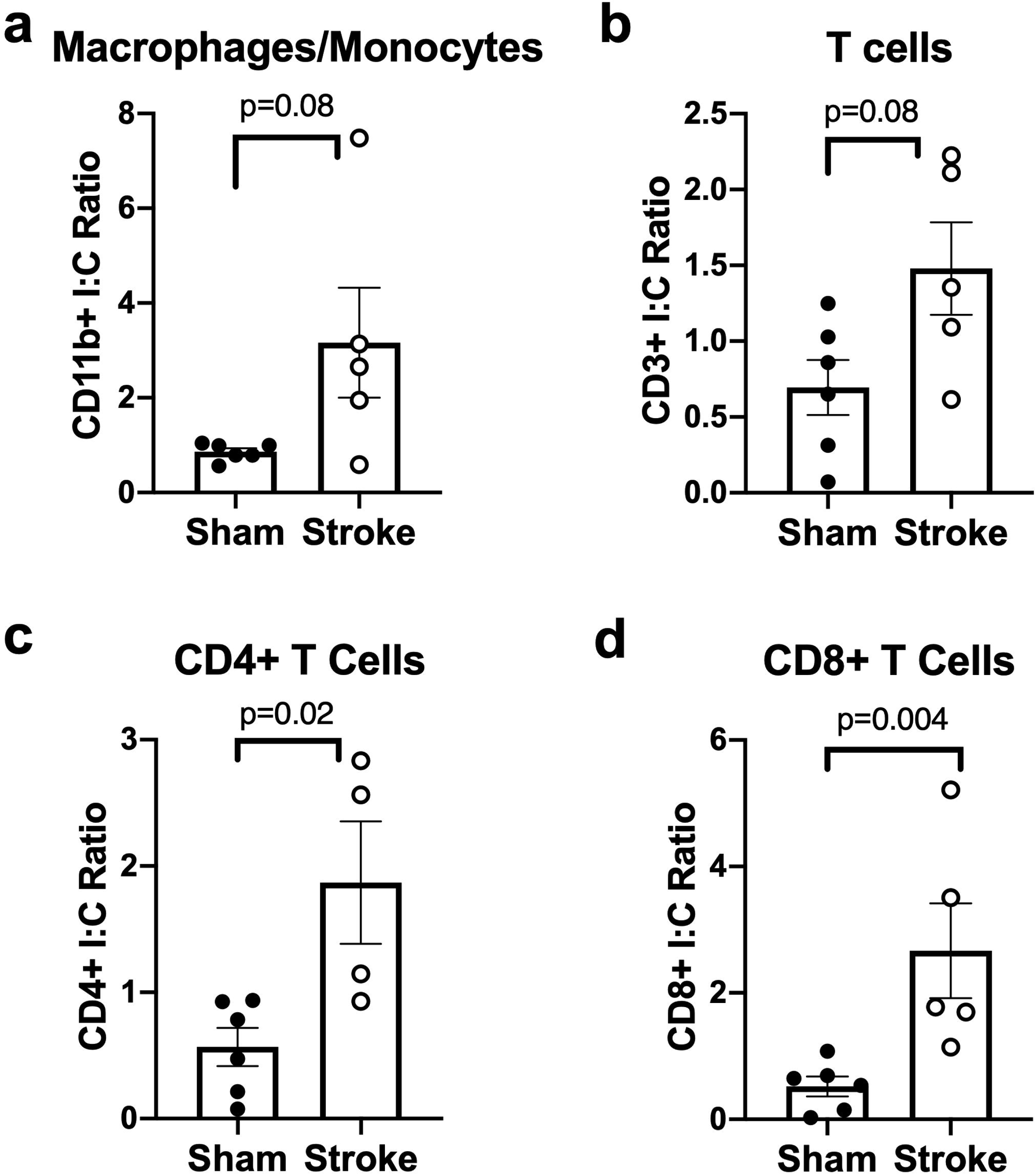
CD4+ and CD8+ T-cell populations are elevated in the ipsilateral cortex at 24 hrs post-stroke. Flow cytometric analysis of murine ipsilateral to contralateral (I:C) hemisphere ratios showed trending or significant increases in (**a**) monocytes/macrophage (p=0.08), (**b**) CD3+ T cells (p=0.08), (**c**) CD4+ T cells (p=0.02), and (**d**) CD8+ T cells (p=0.004) post-stroke compared to sham controls. Data were analyzed using the Mann-Whitney test; sham n=6 and stroke n=4-5.

### Transient MCAO induces acute systemic inflammation

Since stroke induces a systemic proinflammatory response, the serum levels of multiple proinflammatory markers were quantified at 24 hours and 31-36 days post-stroke. P-selectin (**Figure 4a**), vascular endothelial growth factor (VEGF, **Figure 4b**), and monocyte chemoattractant protein 1 (MCP-1/CCL2, **Figure 4c**) were elevated at 24 hours post-stroke and returned to sham levels at 31-36 days post-stroke. Further examination of the endothelial dysfunction markers, E-selectin, L-selectin, vascular cell adhesion protein 1 (VCAM1), and intracellular cell adhesion protein 1 (ICAM1), showed no significant differences between sham and stroke animals at early or late endpoints (**Supplemental Figure 2**). We also observed elevated serum nitric oxide (NO_x_) levels in stroke-injured mice compared to shams at 24 hours post-surgery; however, NO_x_ levels in stroke-injured mice were comparable to shams after 31-36 days (**Figure 4d**). Additionally, we used quantitative RT-PCR analysis to assess differences in liver redox gene expression profiles as previously described [27]. We found that eosinophil peroxidase, interleukin-22, interleukin-6, nitric oxide synthase 2, and tumor necrosis factor-alpha were significantly elevated at 31-36 days post-stroke when normalized to sham controls (**Table 1**).

**Table 1.** Sustained increases in liver pro-inflammatory gene expression post-stroke.

| **Gene** | **Gene Name** | **Assay ID** | **Sham (RQ)** | **Stroke (RQ)** | **Stroke P-value** |
| --- | --- | --- | --- | --- | --- |
| *Epx* | Eosinophil peroxidase | Mm00514768_m1 | 1 | 1.89 | 0.0095 |
| *IL-22* | Interleukin 22 | Mm01226722_g1 | 1 | 1.89 | 0.0095 |
| *IL-6* | Interleukin 6 | Mm00446190_m1 | 1 | 1.73 | 0.035 |
| *Nos2* | Nitric oxide synthase 2, inducible | Mm00440502_m1 | 1 | 1.74 | 0.034 |
| *Tnf* | Tumor necrosis factor $\alpha$ | Mm00443258_m1 | 1 | 1.84 | 0.018 |

Quantitative PCR analysis showed gene expression changes in murine liver tissue post-stroke.
Data were analyzed in DataAssist Software v3.01 using Benjamini-Hochberg False Discovery Rate, where $p < 0.05$ is significant; sham $n=7$ , stroke $n=5$ . RQ = relative quantity (fold change compared to sham).

**Fig. 4.**
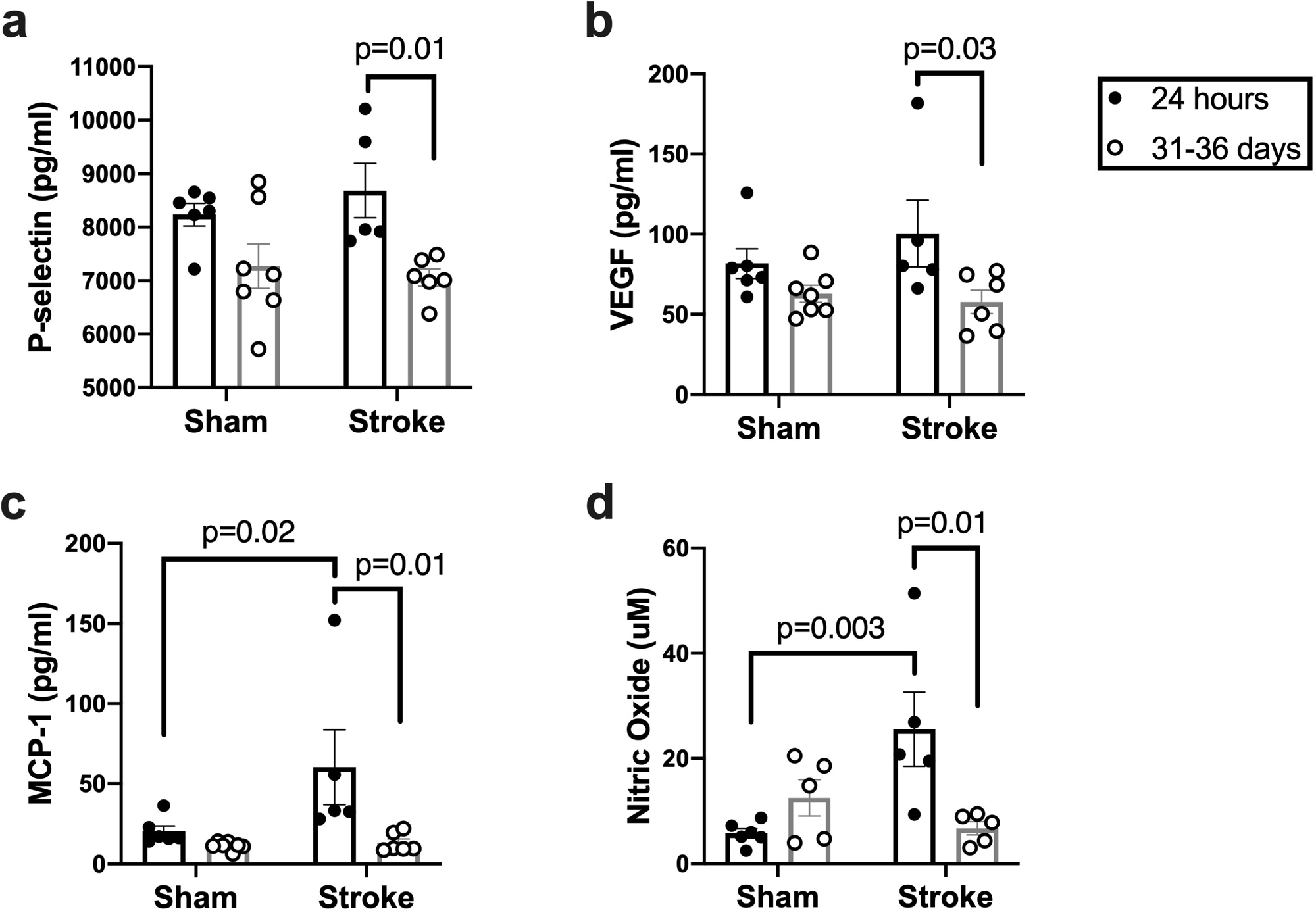
Expression of inflammatory molecules is elevated at 24 hours post-stroke and returns to sham levels by 31-36 days post-stroke. Serum (**a**) P-selectin (p=0.01; main effect of time p=0.001), (**b**) vascular endothelial growth factor (VEGF; p=0.03; main effect of time p=0.01), (**c**) monocyte chemoattractant protein 1 (MCP-1/CCL2; p=0.02; main effect of stroke p=0.01), and (**d**) total nitric oxide (p=0.01, main effect of stroke x time interaction p=0.007), represented as NOx, were elevated at 24 hours post-stroke compared to 31-36 days post-stroke. MCP-1/CCL2 (p=0.02) and nitric oxide (p=0.003) were also significantly increased at 24 hours post-stroke compared to their respective sham controls. Data were analyzed using two-way ANOVA (main effects) followed by Sidak’s multiple comparisons test; n=5-7 per group.

### Transient MCAO induces long-term clinical and behavioral deficits

We next assessed the effect of 60 minutes tMCAO on mortality, clinical scores, and behavioral deficits. First, we observed a 28% mortality level in tMCAO mice throughout the duration of the entire study and did not observe any deaths after 5 days (**Figure 5a**). As expected, stroke-injured mice experienced significant weight loss compared to sham controls during the first week of injury and did not recover to sham levels by the termination of the study (**Figure 5b**). We also found that stroke-injured mice had worse clinical scores compared to sham animals by 24 hours post-injury, and this persisted up to three weeks after stroke (**Figure 5c**). Assessment of spontaneous locomotor activity by open-field testing demonstrate that horizontal movements were significantly decreased in stroke animals compared to sham mice at day 3 post-injury, but recovered to sham levels by day 7 (**Figure 5d**). In contrast, vertical (rearing) movement was impaired in stroke-injured mice for the entire duration of the study (**Figure 5e**). This was mirrored by a diminished level of evoked movement and coordination in stroke mice tested weekly on the rotarod compared to the shams (**Figure 5f**). These results parallel the persistent systemic and neuroinflammation in stroke-injured mice and suggest an interaction between post-stroke inflammation and sustained sensorimotor outcomes.

**Fig. 5.**
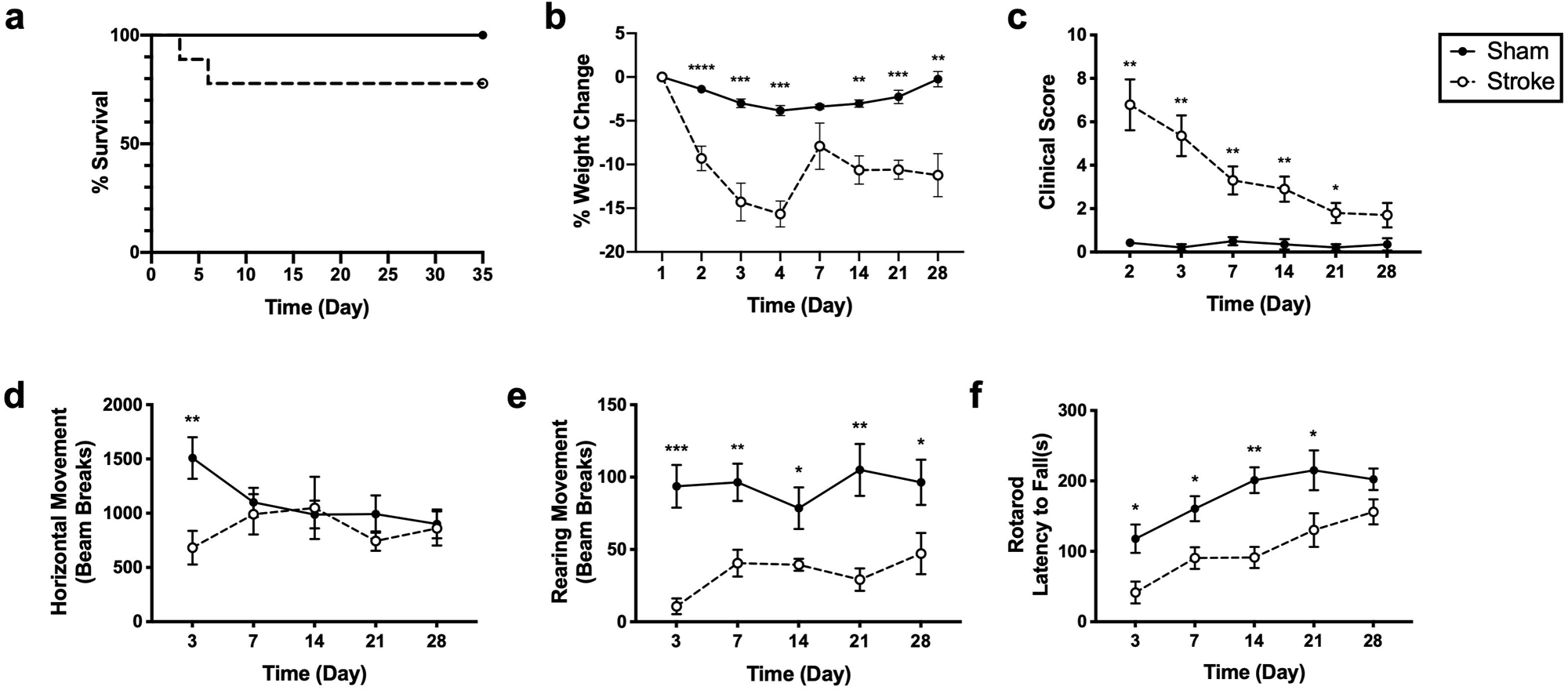
Clinical and behavioral deficits are sustained at 28 days post-stroke. (**a**) Kaplan-Meier curves show higher survival in sham compared to stroke-injured mice. (**b**) Mice with a stroke injury lost more weight than their sham counterparts (main effect of stroke, time, and stroke x time interaction p<0.0001) and (**c**) clinical severity scores remained elevated in stroke-injured mice (main effects of stroke and time p<0.0001, stroke x time interaction p=0.006). Open field testing showed that (**d**) horizontal movement was impaired in the first 3 days post-stroke (main effect of stroke x time interaction p=0.001), while (**e**) rearing movements remained impaired in stroke-injured mice for the duration of the study (main effect of stroke p=0.0009). (**f**) Analysis of evoked movement by rotarod showed motor impairment that persisted up to 21 days post-stroke injury (main effect of stroke p=0.0008 and time p<0.0001). Data were analyzed using repeated measures two-way ANOVA (main effects) with Sidak’s (b, c) or Benjamini, Krieger, and Yekutieli (d, e, f) multiple comparisons tests; n=5-7 per group; *p≤0.05, **p≤0.01, ***p≤0.001, ****p≤0.0001.

### Sustained neuropathology and neuroinflammation post-tMCAO

Histological outcomes were assessed in sham controls and stroke-injured mice to evaluate neuropathological changes at 31-36 days post-surgery. Hematoxylin and eosin staining demonstrated a severe disruption of cortical integrity and a massive infiltration of leukocytes in stroke-injured mice compared to shams (**Figure 6a-b**). Next, we employed immunohistochemical approaches to evaluate the extent of neuroinflammatory changes in the brain post-stroke. The first set of experiments demonstrated an extensive accumulation of CD11b+ immunoreactive cells that was observed in the choroid plexus of mice subjected to tMCAO compared to sham controls (**Figure 6c-d**).

**Fig. 6.**
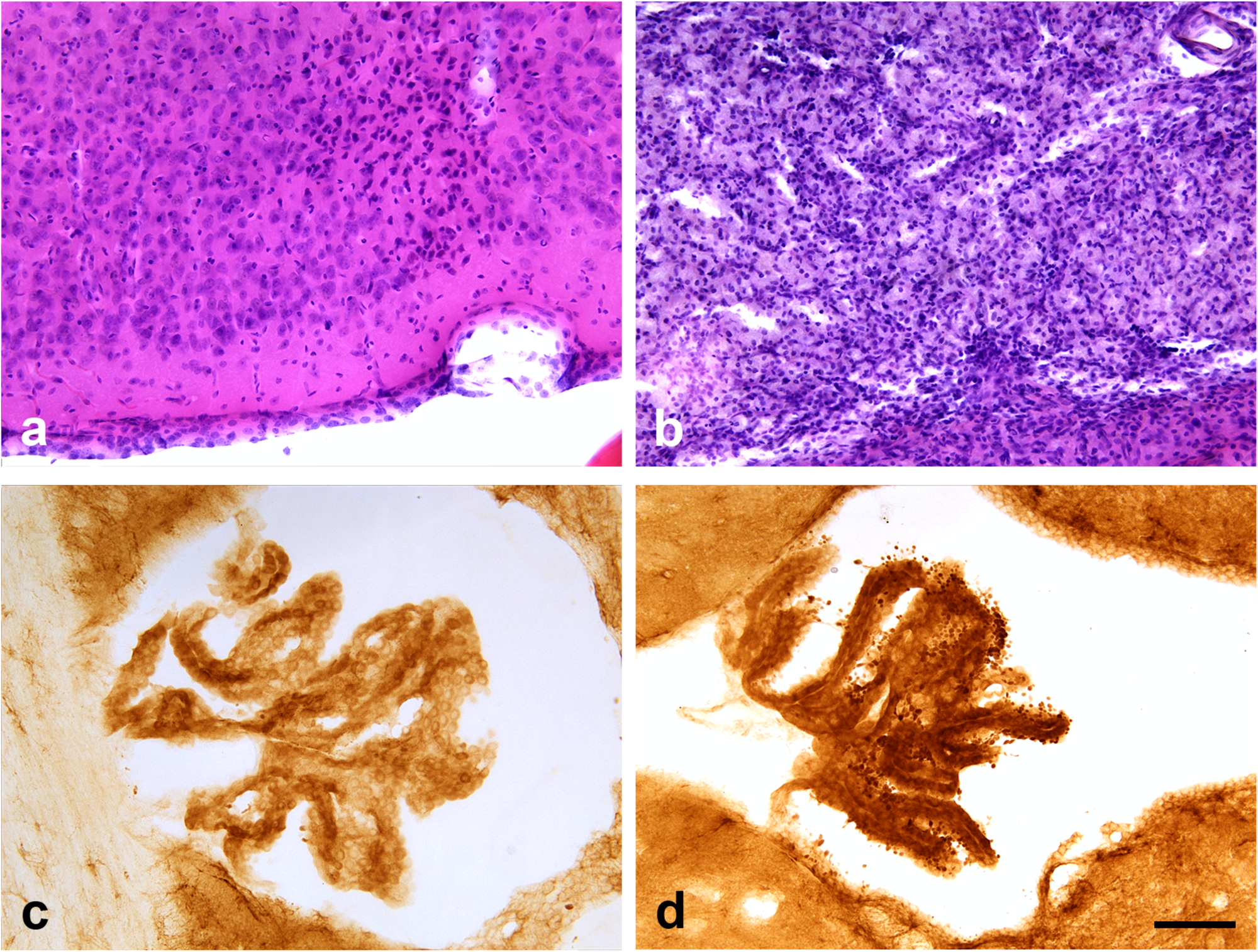
Sustained infiltration of peripheral inflammatory cells 31-36 days post-stroke. Representative hematoxylin and eosin staining in cortex of (**a**) sham compared to (**b**) stroke-injured mice revealed a disruption of cortical integrity and massive influx of inflammatory cells in the penumbra in the stroke-injured mice. Representative immunostaining of CD11b+ cells in choroid plexus shows minimal leukocytes in (**c**) sham controls compared to (**d**) stroke-injured mice. Scale bar = 80 μm.

A second set of studies addressed differences in astrocyte reactivity and microglial activation. Glial fibrillary acidic protein (GFAP) immunostaining in comparable cortical regions from sham controls (**Figure 7a,c**) compared to stroke-injured mice (**Figure 7b,d**) demonstrated elevated immunoreactivity in mice subjected to tMCAO. Higher magnification images from sham-injured mice showed astrocytes with thin processes and faint or incomplete staining of the cell body (**Figure 7b**). In contrast, cortical astrocytes from stroke-injured mice exhibited an activated phenotype as shown by increased GFAP immunoreactivity compared to shams and short, thick cell processes (**Figure 7d**). Quantification of GFAP immunoreactive density demonstrated a significant increase in GFAP staining in stroke-injured mice compared to sham controls (**Figure 7e**). We also evaluated changes in cortical microglia using the microglia/monocyte marker, ionized calcium adapter binding molecule (Iba-1). Sham controls displayed typical microglial morphology in comparable cortical regions (**Figure 8a**) compared to stroke-injured mice, which exhibited dense Iba-1+ immunoreactivity around the infarct (**Figure 8c**). At higher magnification, the morphology of Iba-1+ cells exhibited a generalized surveillance state with light to medium immunostaining of the cell body coupled with multiple long and thin processes (**Figure 8b**). In contrast, experimental stroke resulted in a chronic microglial activation characterized by more extensive Iba-1+ immunoreactivity and shorter, thicker processes that were filled with phagocytic debris presumably from necrotic cells in the core of the infarct (**Figure 8d**). Quantification of the number and area of Iba-1+ cells demonstrated a highly significant increase in stroke-injured mice compared to shams at 31-36 days post-stroke (**Figure 8e-f**).

**Fig. 7.**
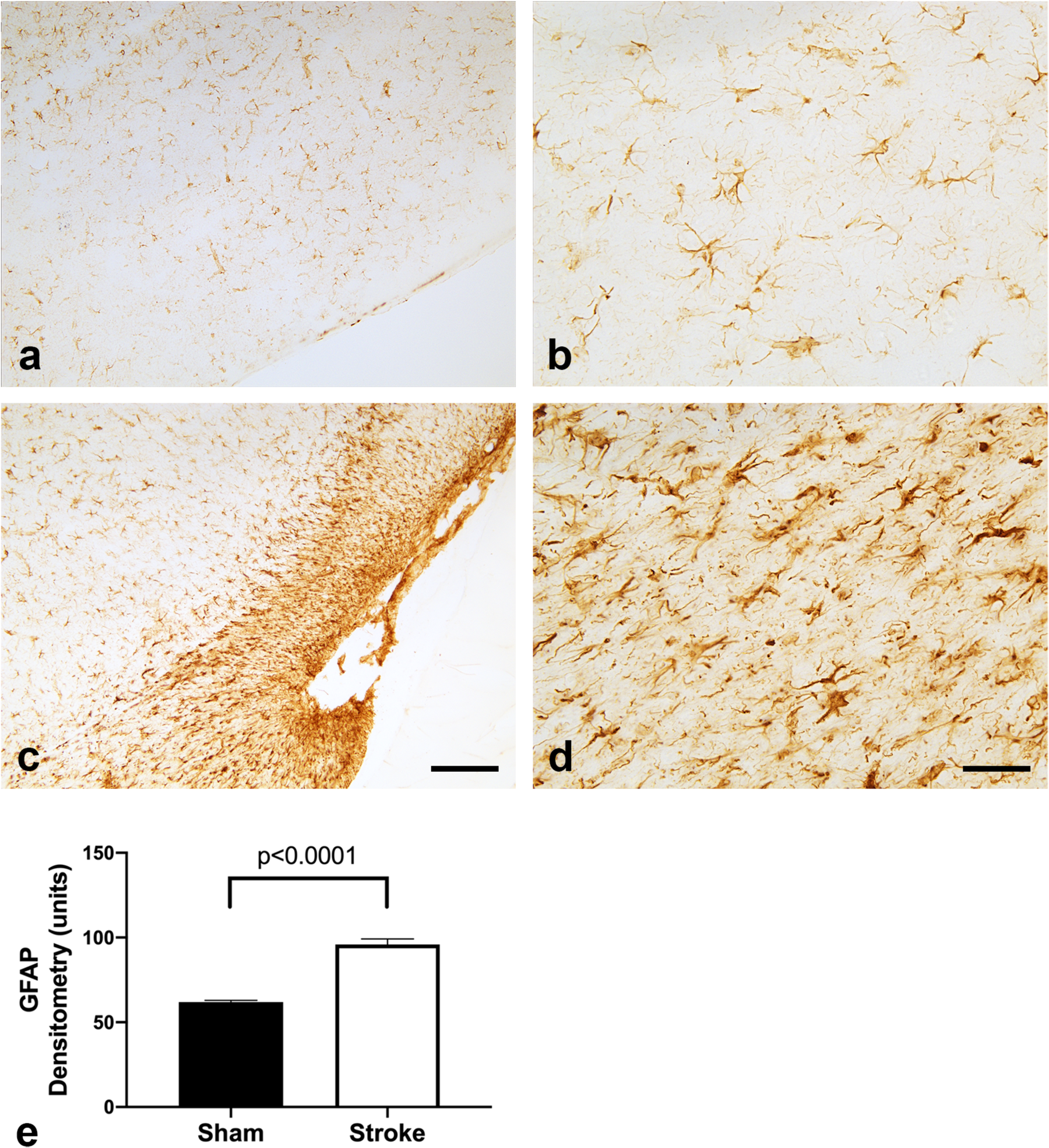
Astrocyte reactivity is elevated at 31-36 days post-stroke. Glial fibrillary acidic protein (GFAP) immunostaining in representative images of cortical astrocytes showed mild immunoreactivity in (**a**) sham controls compared to **(c)** stroke-injured mice. Higher magnification images demonstrate that **(b)** sham surgery was associated with astrocytes with light GFAP immunostaining and long, thin processes, while **(d)** experimental stroke caused dark GFAP immunostaining of astrocytes with short, thick processes. **(e)** Quantification of the density of immunostaining revealed a significant elevation in GFAP in the cortex of stroke animals (p<0.0001); n=4 sham, n=3 stroke animals. Data represent means ± SEM and were analyzed using Student’s t-test. Scale bar in a, c = 160 μm. Scale bar in b, d = 40 μm.

**Fig. 8.**
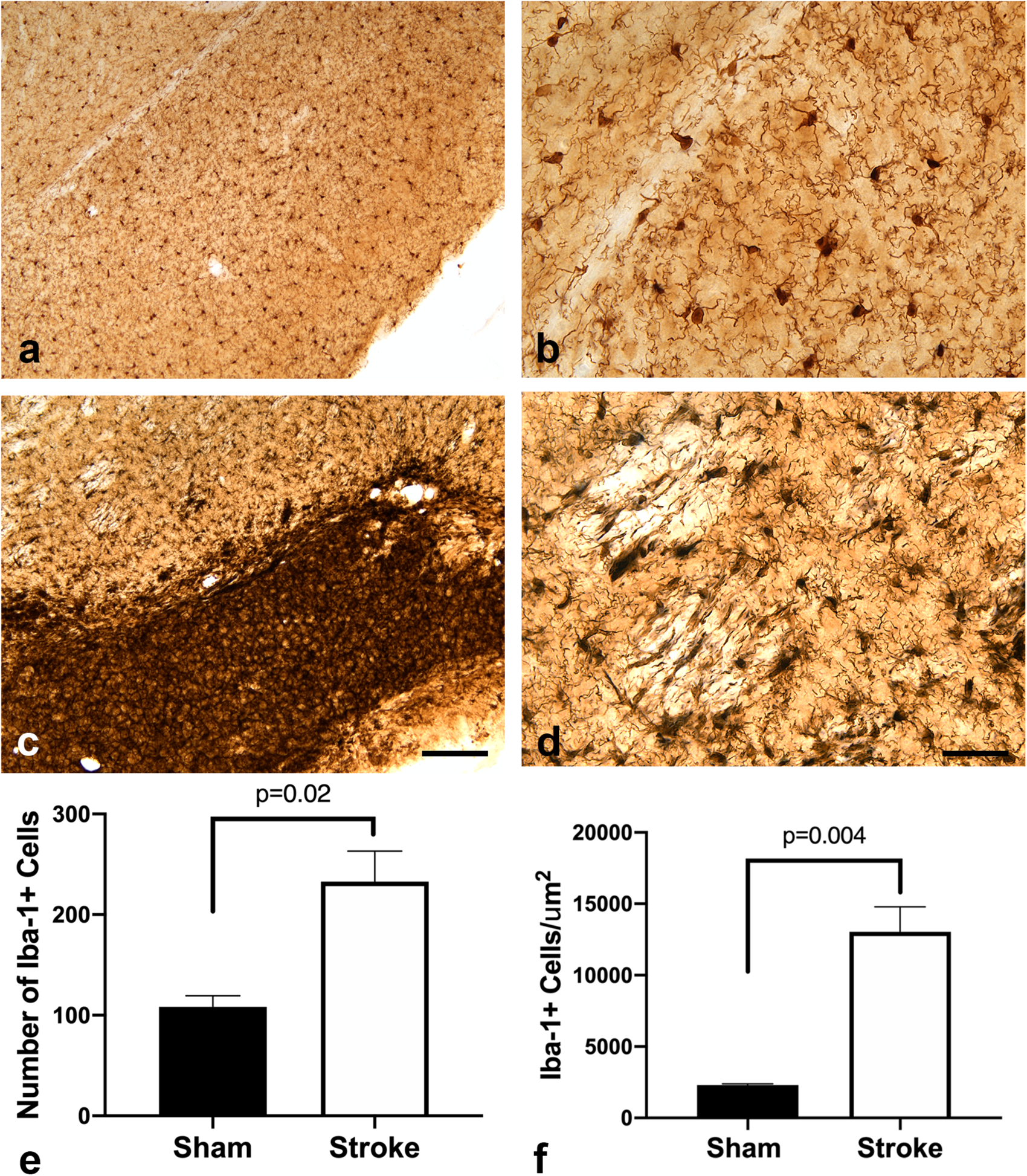
Chronic microglial activation is sustained at 31-36 days following experimental stroke. Microglial/monocyte activation was assessed by ionizing calcium binding adaptor protein -1 (Iba-1) immunoreactivity in cortex. Iba-1 immunostaining in representative cortical images from **(a)** sham and **(c)** stroke-injured mice shows that tMCAO resulted in distinct morphological differences between both groups with a dense accumulation of Iba-1 immunoreactivity in the area of infarct and penumbra. Representative higher magnification images from **(b)** sham mice show Iba-1+ cells with a typical cellular morphology consisting of long, thin processes, while **(d)** Iba-1+ cells in stroke-injured mice show an increased density of immunostaining, an activated morphology with short, thick processes, and accumulation of phagocytic debris. Quantification of Iba-1+ cells showed an increased **(e)** number (p=0.02) and **(f)** area of Iba-1+ cells (p=0.004) in cortical regions of stroke-injured mice. Data were analyzed by Student’s t-test and reported as means ± SEM; n=3 sham, n=2 stroke mice. Scale bar in a, c = 160 μm. Scale bar in b, d = 40 μm.

### Longitudinal disruptions in bacterial diversity and relative abundance after longitudinal tMCAO

The next set of experiments addressed whether long-term changes in the gut microbiota were consistent with observed elevations in brain and systemic inflammation, behavioral deficits, and neuropathology. Fecal pellets were collected from each experimental subjects at baseline (day 0), and on days 3, 14, and 28 after sham or tMCAO surgery followed by 16s rRNA sequencing and phylogenic analysis to uncover taxonomic differences in bacterial abundance. Changes in taxa diversity within respective groups, i.e. sham or stroke, was evaluated by calculating the alpha-diversity. The Shannon alpha-diversity metric showed no longitudinal differences among sham controls (**Figure 9a**); in contrast, this metric was significantly different in stroke-injured mice. We identified significant differences in Shannon-alpha-diversity at 28 days post-stroke compared to baseline and 3 days post-stroke (**Figure 9b**). Next, changes in taxa diversity between respective groups was evaluated by calculating the beta-diversity. Calculation of the Bray-Curtis beta-diversity metric demonstrated significant changes between sham- and stroke-injured mice at 3 days post-stroke, but not at baseline, day 14, or day 28 (**Figure 9c-f**). Taken together, results from our model demonstrate that beta-diversity is altered at earlier time points post-stroke, while alpha-diversity changes occur at later time-points post-stroke.

**Fig. 9.**
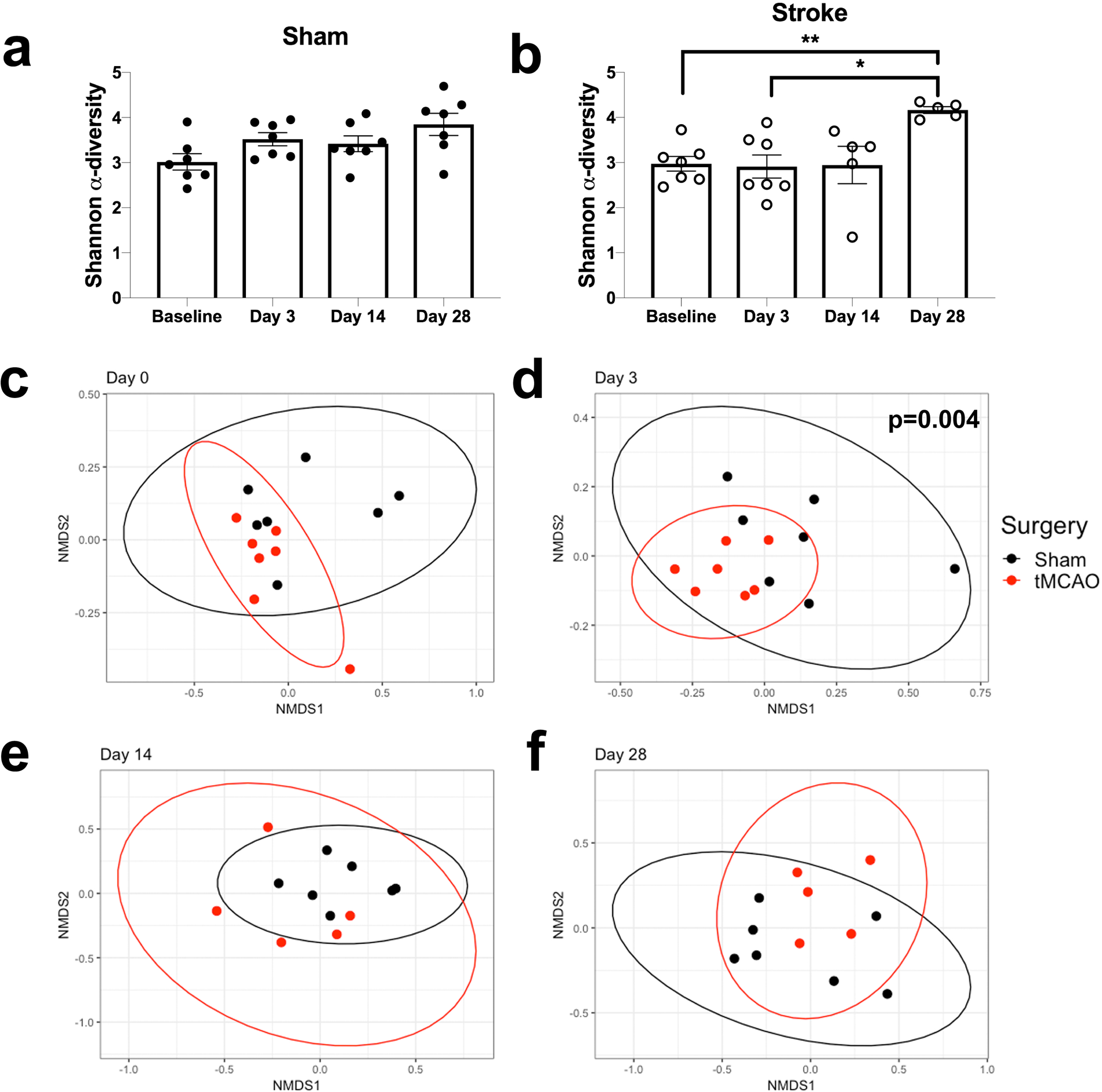
Long-term alterations in alpha-diversity and short-term alterations in beta-diversity occur following ischemic stroke. (**a**). Shannon alpha-diversity showed no significant longitudinal differences in sham mice, while (**b**) significant differences were detected at 28 days post-stroke compared to baseline (**p=0.003) and 3 days post-stroke (*p=0.03). Beta-diversity was not significantly different between sham and stroke-injured mice at **(c)** baseline, while groups exhibited significant dissimilarity at **(d)** 3 days post-stroke (p=0.004); this difference was no longer present by **(e)** 14 days and **(f)** 28 days post-stroke. Alpha-diversity was analyzed using one-way ANOVA (main effect of time p=0.03) with Tukey’s post hoc test. Beta-diversity was calculated using the Bray Curtis distance, visualized using non-metric multidimensional scaling (NMDS), and differences between groups were analyzed using permutation ANOVA (PERMANOVA). Data are reported as means ± SEM ; n=5-7 mice/group.

Taxonomic differences in bacterial abundance were compared to assess longitudinal changes from the phylum to the genus level between sham and stroke-injured mice. We first examined longitudinal changes among bacterial phyla. Pre-surgery results in naïve mice, i.e. baseline results, were not significantly different between surgery groups and were therefore combined; these results showed that Bacteroidetes and Firmicutes comprised the two predominant phyla at baseline, with a larger percentage of Bacteroidetes (**Figure 10a**). We noted an overall effect of surgery on day 3, as Bacteroidetes abundance was decreased in both sham controls and stroke-injured mice along with an increase in Protobacteria abundance among mice with experimental stroke (**Figure 10b**). At day 14 post-stroke, Bacteroidetes abundance in the sham mice recovered to baseline levels, while Bacteroidetes abundance decreased and Firmicutes abundance increased in stroke-injured mice (**Figure 10c**). On day 28, Bacteroidetes abundance remained much lower in stroke-injured mice compared to shams, while Firmicute abundance remained elevated in stroke-injured mice. (**Figure 10d**). The Firmicutes:Bacteroidetes ratio (F:B), where higher values are associated with worse gut health in many gastrointestinal and metabolic disorders, was also calculated. We observed that this ratio is elevated in mice who received a stroke compared to sham controls with a significant interaction between stroke and time and a main effect of stroke (**Figure 10e**).

**Fig. 10.**
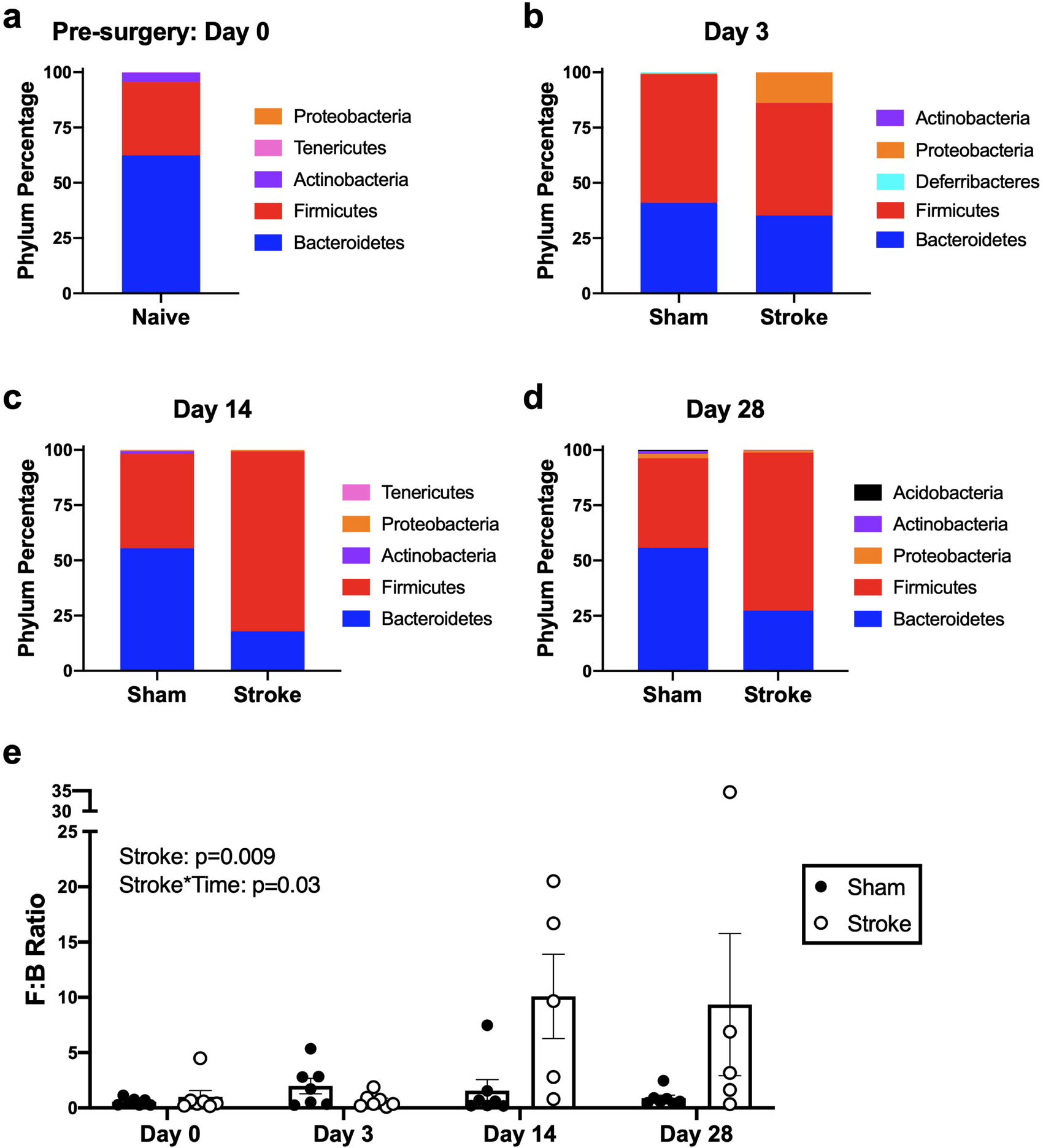
The Firmicutes:Bacteroidetes ratio is elevated at 28 days post-stroke. Relative abundance of the top five bacterial phyla are displayed at day 0 (pre-surgery/baseline) (**a**), day 3 (**b**), day 14 (**c**), and day 28 (**d**) post-tMCAO or sham surgery. Phyla are displayed from most abundant (bottom of graph) to least abundant (top of graph). Calculation of the Firmicutes:Bacteroidetes (F:B) ratio at each time point revealed a stroke x time interaction (p=0.03) and main effect of stroke (p=0.009) that was evident on day 14 and day 28 compared to sham mice (**e**). F:B ratios were analyzed using two-way repeated measures ANOVA with Sidak’s post hoc test. Data are reported as means ± SEM; n=5-7 mice/group.

The longitudinal differences in bacterial abundance between sham- and stroke-injured mice were also evaluated. Since Bacteroidetes and Firmicutes were the two most abundant phyla, we focused on differences among sub-taxa from the phylum level to the family or genus level and reported representative significant or trending results between sham and stroke mice. We found that Firmicutes abundance increased over time in stroke-injured mice and observed significant stroke x time interactions down to the genus level (**Figure 11a**). Concomitant increases in bacterial abundance were observed for the sub-taxa: Clostridia (class), Clostridales (order), *Ruminococcaceae* (family), and *Oscillospira* (genus). In contrast, Bacteroidetes abundance showed a significant decrease or trending decrease over time in stroke-injured mice as evidenced by stroke x time interactions observed down to the family level (**Figure 11b**). Significant and trending decreases were observed for the sub-taxa: Bacteroidia (class), Bacteroidales (order), and *S24-7/Muribaculaceae* (family).Statistical results for the most abundant groups in each taxa as calculated by LMER are summarized in **Supplemental Table 1**.

**Fig. 11.**
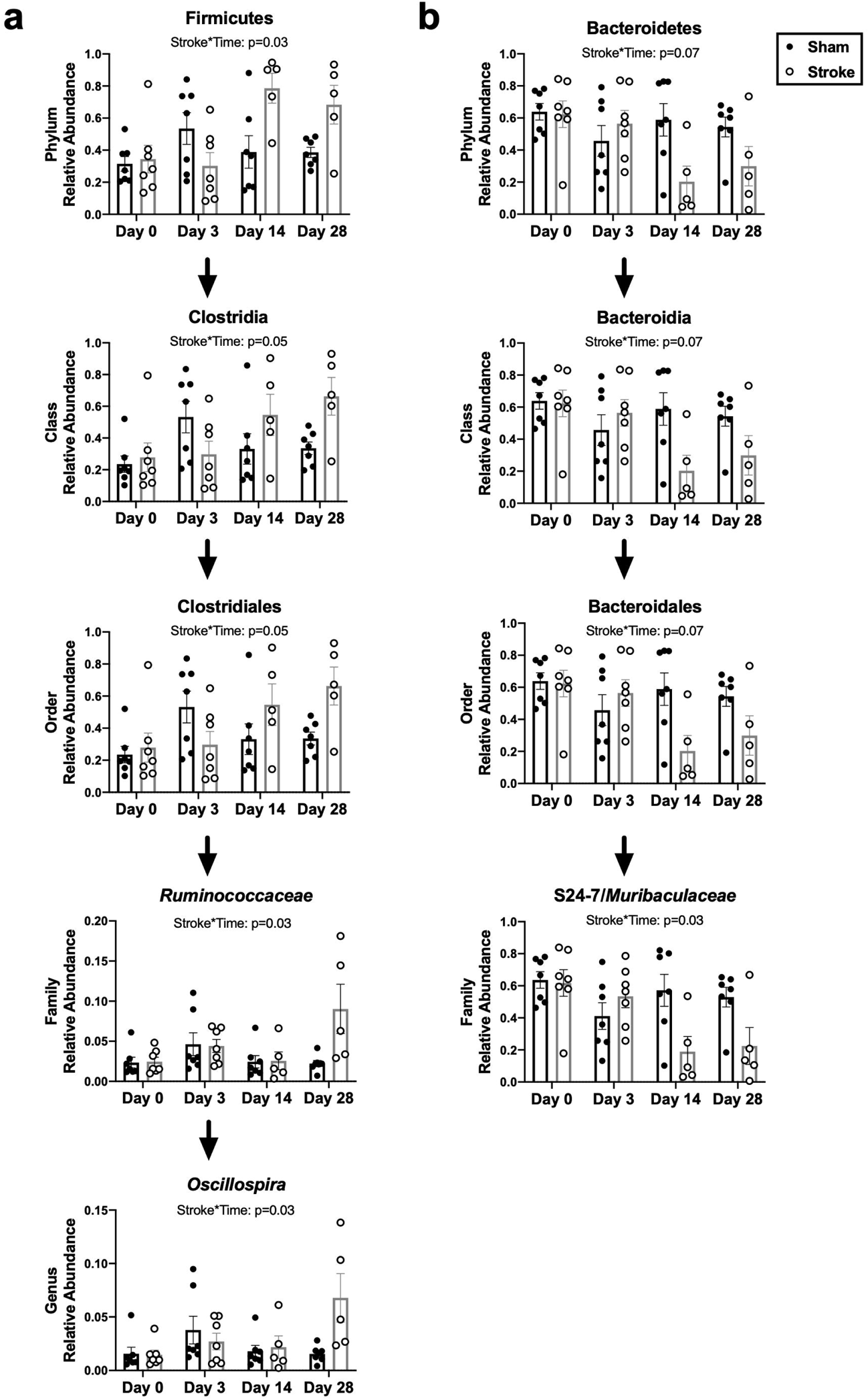
Longitudinal changes in taxa relative abundance 28 days post-stroke show an increase in Firmicutes abundance and decrease in Bacteroidetes abundance in stroke-injured mice. Relative abundance of the most prevalent **(a)** Firmicutes and **(b)** Bacteroidetes bacteria for sham and stroke-injured mice are shown at day 0 (baseline), 3, 14, and 28 days. Significant representative taxa at the class, order, family, and genus levels are shown for each phylum. Data were analyzed by a linear mixed effects model (LMER) and adjusted p-values are reported for each significant stroke x time interaction. Data are represented as means ± SEM; n=5-7 mice/group.

To confirm that our statistical analysis was reproducible, we also analyzed our results using a second method that employs a two-part zero-inflated beta regression model, or the ZIBR model. The ZIBR model is better suited to the analysis of longitudinal datasets but is unable to accommodate for missing observations. Therefore, we did not employ the ZIBR model to report the results in Figure 11. Nevertheless, we used the ZIBR model to perform a comparable analysis and the statistical results are reported in **Supplemental Table 2**. There is relatively strong agreement for results obtained using both the LMER and ZIBR models, which suggests that our results are robust and reproducible.

### Intestinal pathology persists post-tMCAO

In our final set of experiments, we sought to determine whether the differences we observed in bacterial diversity and abundance would be reflected by more severe intestinal pathology in stroke-injured mice compared to sham controls at 31-36 days post-surgery. Periodic acid-Schiff (PAS) staining of the duodenum revealed a thick muscularis mucosae (blue arrow) and a wide crypt depth (black arrow) and moderate infiltration of leukocytes (green arrow) in a representative image from a sham control mouse (**Figure 12 a,c).** In contrast, there is a visible thinning of the muscularis mucosae in stroke-injured mice. The crypt depth is shorter and contains increased numbers of leukocytes compared to shams, and the lamina propria of the villi in stroke mice appear inflamed and edematous (**Figure 12 b,d**).

**Fig. 12.**
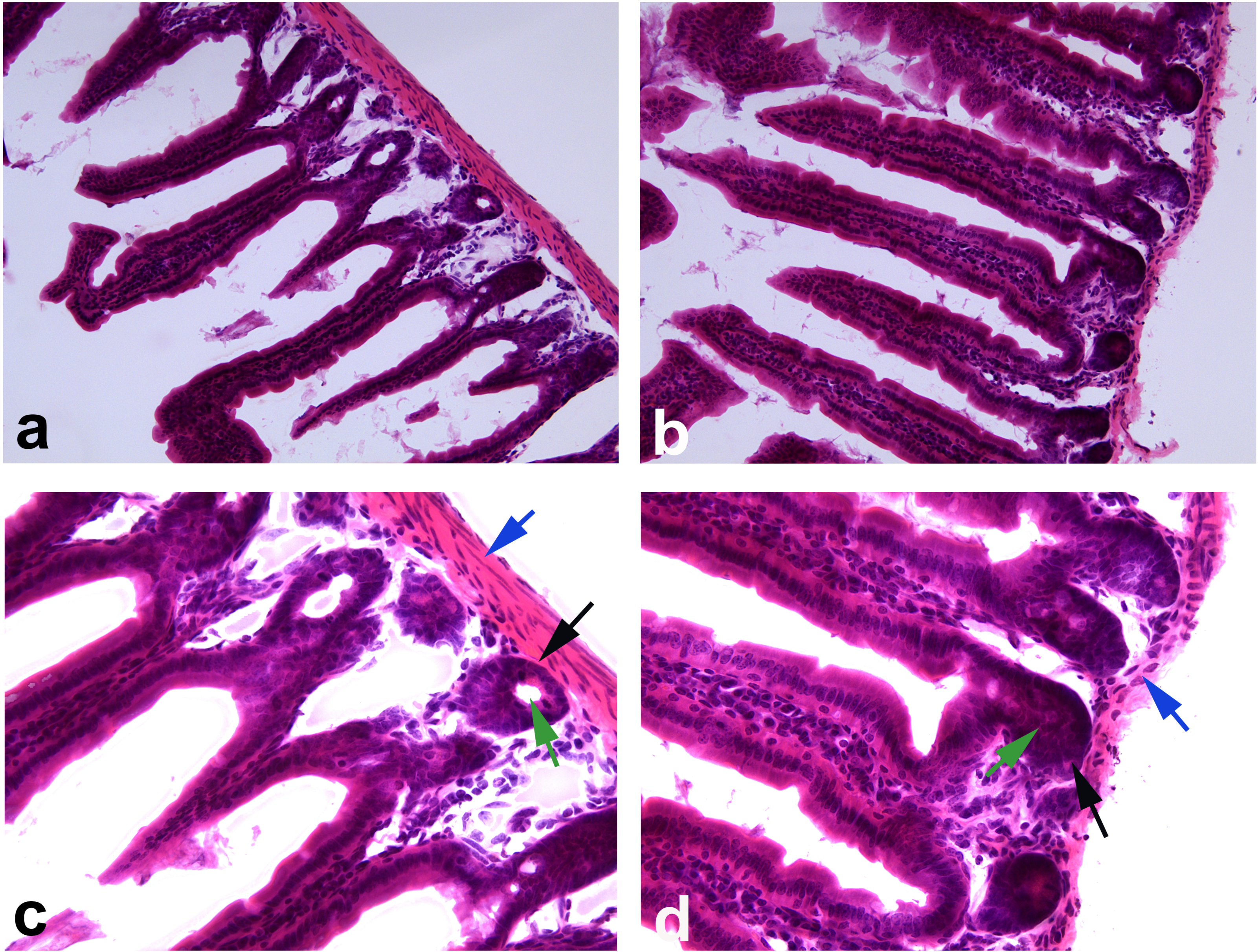
Stroke-injured mice exhibit more severe intestinal pathology compared to sham controls at 31-36 days post-surgery. Representative images of Periodic-Acid Schiff (PAS) staining from (**a,c**) sham control and (**b,d**) stroke-injured mouse duodenum reveals a thicker basement membrane (blue arrows), wider crypt depth (black arrow), and less infiltration of leukocytes (green arrow) in sham compared to stroke intestines. Higher magnification images are shown in (**c**) and (**d**) to illustrate pathology; **a-b**: scale bar = 80 μm; **c-d**: scale bar = 40 μm.

## Discussion

Stroke is an important public health concern, especially within the aging population. In this study, we sought to investigate the long-term changes within the gut-brain-microbiota axis in a transient middle cerebral artery occlusion model of ischemic stroke. We followed a cohort of mice for ∼35 days to examine the natural progression and consequences of ischemic stroke in both the brain and the intestine. First, we demonstrated that there was sustained neuroinflammation and persistent functional deficits at one month post-stroke. Second, sustained alterations in gut microbiota bacterial abundance and intestinal pathology paralleled chronic neuropathological and behavioral deficits. These novel results add to the emerging body of research on stroke and the gut microbiome. Importantly, most prior studies have focused on the early immune changes that alter gut dysbiosis post-stroke. In contrast, our results significantly extend this time frame and clearly demonstrate that post-stroke gut dysbiosis is accompanied by a chronic neuroinflammatory response.

Previous studies have established that acute stroke triggers a number of inflammatory cascades in the brain and systemic immune system [38,39]. Likewise, our transient ischemic stroke model demonstrated an increase in brain CD11b+ monocytes/macrophages and CD3+ T cells, and significant increases in CD4+ T cells and CD8+ T cells by 24 hours post-stroke. Although T cells are typically recruited as part of the adaptive immune response, which is thought to occur several days post-injury, recent work suggests that CD4+ T cells and CD8+ T cells can be recruited within 24 hours post MCAO [40]. Due to the importance of the immune system in stroke, immunomodulatory therapies are under investigation [41]. The depletion of CD4+ T cells and CD8+ T cells in a preclinical tMCAO model has led to significantly reduced infarct volumes 24 hours after stroke, indicating T cells play an important role in neural damage [42]. We also observed that serum levels of the proinflammatory chemokine MCP-1 and nitric oxide (NO) were upregulated 24 hours post-MCAO compared to sham animals. These data confirm that our tMCAO model generated a prototypical inflammatory cascade in both the brain and systemic circulation 24 hours after ischemic injury.

We also observed a prolonged neuroinflammatory response after stroke in our mice at 31-36 days post-injury. This finding is consistent with clinical stroke studies [43–45], yet a precise understanding of the cellular and molecular mechanisms remains unclear. Histological analyses of our stroke brains at 31-36 days post-injury showed evidence of cell death and a glial scar in the cortex. Importantly, we identified clear evidence of microglial/monocyte activation and elevated astrocyte reactivity. Acute cerebral ischemia leads to rapid activation of brain resident microglia, which become phagocytic to promote clearance of debris and cellular repair [46]. The role of microglial cells in stroke is multi-factorial and depends on the type of stroke and timing [46,47]. While there is limited knowledge on the prolonged effects of astrocyte activation post-stroke in humans, GFAP blood concentrations have been associated with stroke severity and long-term functional outcome [48]. The combination of sustained neuroinflammation and systemic immunosuppression leaves stroke patients vulnerable to infection [49]. Therefore, we speculate that it was these prolonged neuro-inflammatory effects that contribute to the functional deficits we observed in our mice post-stroke.

Functional deficits, assessed via sensorimotor function, are one of the most common side effects seen in stroke patients. Up to 70% of stroke survivors experience upper limb (arm and/or hand) paralysis, the most predominant motor impairment after MCA stroke [50,51]. Our tMCAO stroke paradigm demonstrates long-term sensorimotor deficits through voluntary (open-field) and forced (rotarod) movement up to 21 days post-stroke. This is consistent with current literature, which show persistent deficits in both animals and humans [52–55]. Consideration of the long-term consequences of stroke are important because sensorimotor and cognitive deficits can severely affect post-stroke quality of life [56–58]. Our longitudinal study parallels the rodent time frame for functional recovery and extends the translatability of our results to stroke patients. Clinical studies have shown impaired social cognition [59,60], memory deficits [61] and somatosensory deficits [62], which may influence overall functional recovery. Functional recovery is most rapid early after stroke, which is typically considered ≤3 months for humans and ≤1 month for rodents [63]. During this time period the brain should transition from a state of cell death and inflammation to neuronal survival and repair [64]. Yet sustained activation of microglia and astrocytes extends the post-stroke inflammatory dysfunction which is thought to suppress or inhibit neuronal repair mechanisms that are critical for post-stroke functional recovery. Experimental studies using microbiome depletion or manipulation have demonstrated that microbiota composition alone, regardless of an associated disease, can influence behavior [65].

An increasing number of studies have shown that a dysfunctional neuroimmune axis is commonly accompanied by changes in the gut microbiome; however, it is unclear whether these changes are acute and/or chronic. We observed significant differences in beta diversity between sham- and stroke-injured male mice at day 3 post-stroke; this is similar to findings from Singh *et al.*, who have shown substantial changes in gut microbiome beta-diversity 3 days after stroke [13]. Interestingly, our results showed that alpha diversity was significantly increased only in stroke mice on day 28 and not at earlier time points. A study observing fecal microbiome of stroke and transient ischemic attack patients, based on samples taken within 48 hours of hospital admission, showed the patient group had greater diversity than the control group [16]. In addition, a study utilizing fecal microbiota transplantation (FMT) of human stool into rats showed that Shannon alpha diversity was naturally increased 29 and 112 days from the study baseline (day 0) [66]. Therefore, we speculate that stroke-associated gut dysbiosis may contribute to an earlier increase in Shannon alpha diversity compared to sham controls.

Our results also demonstrate an elevated Firmicutes:Bacteroidetes ratio (F:B), a well-established measure of gut dysbiosis, in stroke-injured mice. An increased F:B ratio is most notably associated with metabolic disease, but may help identify gut dysbiosis in a wide range of disorders. Multiple factors influence the F:B ratio, and some of the stroke-associated risk factors known to lead to an increased F:B ratio include poor diet, obesity, and age [67–69]. A recent clinical study demonstrated a decrease in genus *Bacteroides* in fecal samples of stroke patients compared to controls taken within two days of hospital admission [16]. This is consistent with the changes seen in our relative abundance, which showed a depletion of Bacteroidetes in stroke animals by day 3 post-stroke. One of our most interesting findings was the observed decrease in abundance of the S24-7 family. Although this taxon has been frequently identified in non-primate microbiome samples, little is known about S24-7, recently renamed *Muribaculaceae* [70]. Although *Muribaculaceae* is presumed to be one of the most-prevalent commensal taxa in mammals [71], it has only recently been cultured and thus, genomic studies are limited [72]. A recent study has suggested that *Muribaculaceae* may fulfill a set of similar duties in mice as *Bacteroides* do in humans, based on the fermentation pathways identified within its metagenome [73].

The observation that bacterial abundance of Firmicutes were elevated during stroke is consistent with the emerging concept that deleterious bacteria are elevated post-stroke and may contribute to poor stroke recovery. We found that *Ruminococcaceae* exhibited a significant stroke x time interaction that was most evident at day 28 post-stroke. Interestingly, changes in some *Ruminococcaceae* have been associated with behavioral deficits and social avoidance in mice [74,75]. Human fecal samples taken from stroke patients within 48 hours of hospital admission showed an increase in *Ruminococcaceae*_UCG_005; and three genera of *Ruminococcaceae* were consistently upregulated in severe stroke compared to mild stroke patients [76]. The abundance of *Ruminococcaceae* in neurocritically ill patients has been negatively associated with length of hospital stay [77]. Additionally, the abundance of *Oscillospira*, the most abundant genus we identified in the family *Ruminococcaceae*, has been shown to increase with age; this is consistent with an increased risk of stroke in aged individuals [78].

As many as 50% of stroke survivors also experience GI complications, which results in a higher risk of mortality and worsened neurologic outcomes in these patients [3,4]. While several studies have revealed damage to the intestinal mucosa soon after experimental stroke [79,80], fewer studies have shown long-term effects. We observed that prolonged intestinal inflammatory pathology accompanies the changes that we see in bacterial abundance. More recently, mucosal and villi damage has been observed in the colons of cynomolgus monkeys at 1.5, 6, and 12 months after permanent MCAO [81]. Although specific mechanisms linking gut dysbiosis and stroke remain elusive, the most likely mechanisms include: increased permeability of the intestinal barrier and bacterial motility, suppression of systemic immunity, release of proinflammatory factors from the brain infarct, and activation of the sympathetic nervous system [82,83].

Our results, as well as the results of others in the field, suggest that manipulation of the gut microbiota may benefit stroke recovery and improve long-term outcomes. Studies using probiotic treatment with *Lactobacillus plantarum* ZDY2013 [84], *Clostridium butyricum* [85], or probiotic mixtures [86,87] have shown improved outcomes in a number of stroke-related disorders. Mediation of bacterial abundances likely affect the percentage of available metabolites, primarily short-chain fatty acids (SCFA; i.e., acetate, propionate, and butyrate) that are produced in the gut; all three of these metabolites have been shown to have numerous beneficial effects on human health [88,89]. Stroke patients have been shown to have altered levels of SCFA in their stool compared to controls [90]. Pre-clinical studies have shown that SCFA supplementation improves motor function [91], reduced neurological impairment, and reduced cerebral infarct volume [92] post-stroke. In more severe cases where supplementation with probiotics or bacterial metabolites is inadequate, it has been suggested that total repopulation of the intestinal tract may be possible using FMT. Although it is best known for its clinical applications in patients with recurrent *Clostridium difficile* infection, FMT has yielded improved post-stroke outcomes in preclinical studies [14] and results may last for several weeks as demonstrated in models of colitis [66].

In conclusion, the microbiome is an increasingly attractive target for a number of neurological and neuroinflammatory diseases, including stroke. We have shown that ischemic stroke results in persistent behavioral deficits, prolonged brain and intestinal inflammation, and continual dysbiosis of the gut microbiota. We have discussed several methods of microbiota manipulation, including the use of probiotics or supplements, SCFAs, and FMT, as a way to replenish the abundance of bacteria taxa that are disrupted after stroke. While therapeutic strategies to target the gut microbiome are in their infancy, current efforts are underway to incorporate targeted modulation of gut microbiome in personalized medicine [93]. Taken together, this study provides novel insights into the longitudinal changes that occur within the microbiota-gut-brain axis in ischemic stroke. Our results establish a foundation for future studies to examine how the manipulation of gut microbiome and GI function may improve both immune function and functional recovery post-stroke.

## Acknowledgments

We wish to thank Dr. Eric Kelley for the use of his Sievers NOA 208 instrument.

## Figure Legends

**Supplemental Figure 1 Brain infiltration of monocytes, neutrophils and B-cells is not significantly different between sham controls and stroke-injured mice at 24 hours post-stroke.** Flow cytometric analysis of rodent ipsilateral to contralateral (I:C) hemisphere ratios between sham and stroke-injured mice showed no differences between **(a)** Ly6C+ monocytes, **(b)** Ly6G+ neutrophils, or **(c)** CD19+ B cells. Data were analyzed by Mann-Whitney test and represent mean ± SEM; sham n=6 and stroke n=5.

**Supplemental Figure 2 Serum levels of proinflammatory markers with no difference between sham controls and stroke-injured mice.** No significant differences between serum levels of (**a**) E-selectin, (**b**) L-selectin, (**c**) TNF-α, (**d**) VCAM-1, or (**e**) ICAM-1 were detected between sham and stroke mice at 24 hours or 31-36 days post-surgery. Data were analyzed using two-way repeated measures ANOVA and represent mean ± SEM; n=5-7 mice/group.

## References

1. Schaller BJ, Graf R, Jacobs AH. Pathophysiological changes of the gastrointestinal tract in ischemic stroke. Am J Gastroenterol. 2006;101:1655–65.

2. Harari D, Norton C, Lockwood L, Swift C. Treatment of Constipation and Fecal Incontinence in Stroke Patients: Randomized Controlled Trial. Stroke. 2004;35:2549–55.

3. Camara-Lemarroy CR, Ibarra-Yruegas BE, Gongora-Rivera F. Gastrointestinal complications after ischemic stroke. Journal of the Neurological Sciences. 2014;346:20–5.

4. Wen SW, Wong CHY. An unexplored brain-gut microbiota axis in stroke. Gut Microbes. 2017;8:601–6.

5. Dinan TG, Cryan JF. The Microbiome-Gut-Brain Axis in Health and Disease. Gastroenterology Clinics of North America. 2017;46:77–89.

6. Winek K, Dirnagl U, Meisel A. The Gut Microbiome as Therapeutic Target in Central Nervous System Diseases: Implications for Stroke. Neurotherapeutics. 2016;13:762–74.

7. Tremlett H, Bauer KC, Appel-Cresswell S, Finlay BB, Waubant E. The gut microbiome in human neurological disease: A review. Ann Neurol. 2017;81:369–82.

8. Burokas A, Moloney RD, Dinan TG, Cryan JF. Microbiota Regulation of the Mammalian Gut–Brain Axis. Advances in Applied Microbiology [Internet]. Elsevier; 2015 [cited 2019 Mar 19]. p. 1–62. Available from: https://linkinghub.elsevier.com/retrieve/pii/S0065216415000027

9. Wang Y, Kasper LH. The role of microbiome in central nervous system disorders. Brain, Behavior, and Immunity. 2014;38:1–12.

10. Das R, Kanungo MS. Effects of polyamines on in vitro phosphorylation and acetylation of histones of the cerebral cortex of rats of various ages. Biochem Biophys Res Commun. 1979;90:708–14.

11. Kelly JR, Borre Y, O’ Brien C, Patterson E, El Aidy S, Deane J, et al. Transferring the blues: Depression-associated gut microbiota induces neurobehavioural changes in the rat. Journal of Psychiatric Research. 2016;82:109–18.

12. Sherwin E, Dinan TG, Cryan JF. Recent developments in understanding the role of the gut microbiota in brain health and disease: The gut microbiota in brain health and disease. Annals of the New York Academy of Sciences. 2018;1420:5–25.

13. Singh V, Roth S, Llovera G, Sadler R, Garzetti D, Stecher B, et al. Microbiota Dysbiosis Controls the Neuroinflammatory Response after Stroke. Journal of Neuroscience. 2016;36:7428–40.

14. Spychala MS, Venna VR, Jandzinski M, Doran SJ, Durgan DJ, Ganesh BP, et al. Age-related changes in the gut microbiota influence systemic inflammation and stroke outcome: Age-Related Changes in the Gut Microbiota. Annals of Neurology [Internet]. 2018 [cited 2018 Aug 30]; Available from: http://doi.wiley.com/10.1002/ana.25250

15. Stanley D, Moore RJ, Wong CHY. An insight into intestinal mucosal microbiota disruption after stroke. Scientific Reports [Internet]. 2018 [cited 2018 May 24];8. Available from: http://www.nature.com/articles/s41598-017-18904-8

16. Yin J, Liao S, He Y, Wang S, Xia G, Liu F, et al. Dysbiosis of Gut Microbiota With Reduced Trimethylamine□N□Oxide Level in Patients With Large□Artery Atherosclerotic Stroke or Transient Ischemic Attack. Journal of the American Heart Association. 2015;4:e002699.

17. Zeng X, Gao X, Peng Y, Wu Q, Zhu J, Tan C, et al. Higher Risk of Stroke Is Correlated With Increased Opportunistic Pathogen Load and Reduced Levels of Butyrate-Producing Bacteria in the Gut. Frontiers in Cellular and Infection Microbiology [Internet]. 2019 [cited 2019 Mar 27];9. Available from: https://www.frontiersin.org/article/10.3389/fcimb.2019.00004/full

18. Benakis C, Brea D, Caballero S, Faraco G, Moore J, Murphy M, et al. Commensal microbiota affects ischemic stroke outcome by regulating intestinal γΔ T cells. Nature Medicine. 2016;22:516–23.

19. Adnan S, Nelson JW, Ajami NJ, Venna VR, Petrosino JF, Bryan RM, et al. Alterations in the gut microbiota can elicit hypertension in rats. Physiological Genomics. 2017;49:96–104.

20. Sert NP du, Hurst V, Ahluwalia A, Alam S, Altman DG, Avey MT, et al. Revision of the ARRIVE guidelines: rationale and scope. BMJ Open Science. 2018;2:e000002.

21. Liebeskind DS, Derdeyn CP, Wechsler LR, for the STAIR X Consortium*, Albers G, Ankerud EP, et al. STAIR X: Emerging Considerations in Developing and Evaluating New Stroke Therapies. Stroke. 2018;49:2241–7.

22. Doll DN, Hu H, Sun J, Lewis SE, Simpkins JW, Ren X. Mitochondrial crisis in cerebrovascular endothelial cells opens the blood-brain barrier. Stroke. 2015;46:1681–9.

23. Chen J, Zhang C, Jiang H, Li Y, Zhang L, Robin A, et al. Atorvastatin Induction of VEGF and BDNF Promotes Brain Plasticity after Stroke in Mice. Journal of Cerebral Blood Flow & Metabolism. 2005;25:281–90.

24. Frank MG, Wieseler-Frank JL, Watkins LR, Maier SF. Rapid isolation of highly enriched and quiescent microglia from adult rat hippocampus: Immunophenotypic and functional characteristics. Journal of Neuroscience Methods. 2006;151:121–30.

25. Mandler WK, Nurkiewicz TR, Porter DW, Kelley EE, Olfert IM. Microvascular Dysfunction Following Multiwalled Carbon Nanotube Exposure Is Mediated by Thrombospondin-1 Receptor CD47. Toxicol Sci. 2018;165:90–9.

26. Brown CM, Xu Q, Okhubo N, Vitek MP, Colton CA. Androgen-Mediated Immune Function Is Altered by the Apolipoprotein E Gene. Endocrinology. Oxford Academic; 2007;148:3383–90.

27. Hoos MD, Vitek MP, Ridnour LA, Wilson J, Jansen M, Everhart A, et al. The impact of human and mouse differences in NOS2 gene expression on the brain’s redox and immune environment. Mol Neurodegener. 2014;9:50.

28. Brichacek AL, Benkovic SA, Chakraborty S, Nwafor DC, Wang W, Jun S, et al. Systemic inhibition of tissue-nonspecific alkaline phosphatase alters the brain-immune axis in experimental sepsis. Sci Rep. 2019;9:18788.

29. Nwafor DC, Chakraborty S, Brichacek AL, Jun S, Gambill CA, Wang W, et al. Loss of tissue-nonspecific alkaline phosphatase (TNAP) enzyme activity in cerebral microvessels is coupled to persistent neuroinflammation and behavioral deficits in late sepsis. Brain, Behavior, and Immunity. 2020;84:115–31.

30. Fadrosh DW, Ma B, Gajer P, Sengamalay N, Ott S, Brotman RM, et al. An improved dual-indexing approach for multiplexed 16S rRNA gene sequencing on the Illumina MiSeq platform. Microbiome. 2014;2:6.

31. Callahan BJ, McMurdie PJ, Rosen MJ, Han AW, Johnson AJA, Holmes SP. DADA2: High resolution sample inference from Illumina amplicon data. Nat Methods. 2016;13:581–3.

32. DeSantis TZ, Hugenholtz P, Larsen N, Rojas M, Brodie EL, Keller K, et al. Greengenes, a Chimera-Checked 16S rRNA Gene Database and Workbench Compatible with ARB. Appl Environ Microbiol. 2006;72:5069–72.

33. Oksanen J, Guillaume Blanchet F, Friendly M, Kindt R, Legendre P, McGlinn D, et al. vegan: Community Ecology Package [Internet]. 2018. Available from: https://CRAN.R-project.org/package=vegan

34. McMurdie PJ, Holmes S. Waste Not, Want Not: Why Rarefying Microbiome Data Is Inadmissible. McHardy AC, editor. PLoS Computational Biology. 2014;10:e1003531.

35. Bates D, Mächler M, Bolker B, Walker S. Fitting Linear Mixed-Effects Models Using lme4. Journal of Statistical Software [Internet]. 2015 [cited 2019 Aug 12];67. Available from: http://www.jstatsoft.org/v67/i01/

36. Chen EZ, Li H. A two-part mixed-effects model for analyzing longitudinal microbiome compositional data. Bioinformatics. 2016;32:2611–7.

37. R Core Team. R: A language and environment for statistical computing. R Foundation for Statistical Computing, Vienna, Austria; 2019.

38. Chamorro Á, Meisel A, Planas AM, Urra X, van de Beek D, Veltkamp R. The immunology of acute stroke. Nature Reviews Neurology. 2012;8:401–10.

39. Iadecola C, Anrather J. The immunology of stroke: from mechanisms to translation. Nature Medicine. 2011;17:796–808.

40. Chu HX, Kim HA, Lee S, Moore JP, Chan CT, Vinh A, et al. Immune Cell Infiltration in Malignant Middle Cerebral Artery Infarction: Comparison with Transient Cerebral Ischemia. Journal of Cerebral Blood Flow & Metabolism. 2014;34:450–9.

41. Malone K, Amu S, Moore AC, Waeber C. Immunomodulatory Therapeutic Strategies in Stroke. Front Pharmacol. 2019;10:630.

42. Yilmaz G, Arumugam TV, Stokes KY, Granger DN. Role of T Lymphocytes and Interferon-γ in Ischemic Stroke. Circulation. 2006;113:2105–12.

43. Vidale S, Consoli A, Arnaboldi M, Consoli D. Postischemic Inflammation in Acute Stroke. J Clin Neurol. 2017;13:1.

44. Fang M, Zhong L, Jin X, Cui R, Yang W, Gao S, et al. Effect of Inflammation on the Process of Stroke Rehabilitation and Poststroke Depression. Front Psychiatry. 2019;10:184.

45. Anrather J, Iadecola C. Inflammation and Stroke: An Overview. Neurotherapeutics. 2016;13:661–70.

46. Benakis C, Garcia-Bonilla L, Iadecola C, Anrather J. The role of microglia and myeloid immune cells in acute cerebral ischemia. Front Cell Neurosci [Internet]. 2015 [cited 2020 Apr 6];8. Available from: http://journal.frontiersin.org/article/10.3389/fncel.2014.00461/abstract

47. Kanazawa M, Ninomiya I, Hatakeyama M, Takahashi T, Shimohata T. Microglia and Monocytes/Macrophages Polarization Reveal Novel Therapeutic Mechanism against Stroke. IJMS. 2017;18:2135.

48. Wunderlich MT, Wallesch CW, Goertler M. Release of glial fibrillary acidic protein is related to the neurovascular status in acute ischemic stroke. Eur J Neurol. England; 2006;13:1118–23.

49. Boaden E, Lyons M, Singhrao SK, Dickinson H, Leathley M, Lightbody CE, et al. Oral flora in acute stroke patients: A prospective exploratory observational study. Gerodontology. 2017;34:343–56.

50. Borschmann KN, Hayward KS. Recovery of upper limb function is greatest early after stroke but does continue to improve during the chronic phase: a two-year, observational study. Physiotherapy. 2020;107:216–23.

51. Lang CE, Schieber MH. Differential Impairment of Individuated Finger Movements in Humans After Damage to the Motor Cortex or the Corticospinal Tract. Journal of Neurophysiology. 2003;90:1160–70.

52. Pin-Barre C, Laurin J. Physical Exercise as a Diagnostic, Rehabilitation, and Preventive Tool: Influence on Neuroplasticity and Motor Recovery after Stroke. Neural Plasticity. 2015;2015:1–12.

53. Hatem SM, Saussez G, della Faille M, Prist V, Zhang X, Dispa D, et al. Rehabilitation of Motor Function after Stroke: A Multiple Systematic Review Focused on Techniques to Stimulate Upper Extremity Recovery. Front Hum Neurosci [Internet]. 2016 [cited 2020 Apr 6];10. Available from: http://journal.frontiersin.org/Article/10.3389/fnhum.2016.00442/abstract

54. Balkaya M, Kröber JM, Rex A, Endres M. Assessing Post-Stroke Behavior in Mouse Models of Focal Ischemia. J Cereb Blood Flow Metab. 2013;33:330–8.

55. Balkaya MG, Trueman RC, Boltze J, Corbett D, Jolkkonen J. Behavioral outcome measures to improve experimental stroke research. Behav Brain Res. Netherlands; 2018;352:161–71.

56. Bansil S, Prakash N, Kaye J, Wrigley S, Manata C, Stevens-Haas C, et al. Movement disorders after stroke in adults: a review. Tremor Other Hyperkinet Mov (N Y). 2012;2.

57. Nakawah MO, Lai EC. Post-stroke dyskinesias. Neuropsychiatric Disease and Treatment. 2016;Volume 12:2885–93.

58. Kalaria RN, Akinyemi R, Ihara M. Stroke injury, cognitive impairment and vascular dementia. Biochim Biophys Acta. 2016/01/22. Elsevier Pub. Co; 2016;1862:915–25.

59. Nijsse B, Spikman JM, Visser-Meily JM, de Kort PL, van Heugten CM. Social Cognition Impairments in the Long Term Post Stroke. Archives of Physical Medicine and Rehabilitation [Internet]. 2019 [cited 2019 May 1]; Available from: https://linkinghub.elsevier.com/retrieve/pii/S0003999319301492

60. Nijsse B, Spikman JM, Visser-Meily JMA, de Kort PLM, van Heugten CM. Social cognition impairments are associated with behavioural changes in the long term after stroke. Pavlova MA, editor. PLoS ONE. 2019;14:e0213725.

61. das Nair R, Cogger H, Worthington E, Lincoln NB. Cognitive rehabilitation for memory deficits after stroke. Cochrane Stroke Group, editor. Cochrane Database of Systematic Reviews [Internet]. 2016 [cited 2020 Apr 6]; Available from: http://doi.wiley.com/10.1002/14651858.CD002293.pub3

62. Kessner SS, Bingel U, Thomalla G. Somatosensory deficits after stroke: a scoping review. Topics in Stroke Rehabilitation. 2016;23:136–46.

63. Caleo M. Rehabilitation and plasticity following stroke: Insights from rodent models. Neuroscience. 2015;311:180–94.

64. Cramer SC. Repairing the human brain after stroke: I. Mechanisms of spontaneous recovery. Ann Neurol. United States; 2008;63:272–87.

65. Vuong HE, Yano JM, Fung TC, Hsiao EY. The Microbiome and Host Behavior. Annual Review of Neuroscience. 2017;40:21–49.

66. Lleal M, Sarrabayrouse G, Willamil J, Santiago A, Pozuelo M, Manichanh C. A single faecal microbiota transplantation modulates the microbiome and improves clinical manifestations in a rat model of colitis. EBioMedicine. 2019;48:630–41.

67. Ley RE, Turnbaugh PJ, Klein S, Gordon JI. Microbial ecology: human gut microbes associated with obesity. Nature. 2006;444:1022–3.

68. Voreades N, Kozil A, Weir TL. Diet and the development of the human intestinal microbiome. Frontiers in Microbiology [Internet]. 2014 [cited 2019 Apr 21];5. Available from: http://journal.frontiersin.org/article/10.3389/fmicb.2014.00494/abstract

69. Mariat D, Firmesse O, Levenez F, Guimarăes V, Sokol H, Doré J, et al. The Firmicutes/Bacteroidetes ratio of the human microbiota changes with age. BMC Microbiology. 2009;9:123.

70. Lagkouvardos I, Lesker TR, Hitch TCA, GÁlvez EJC, Smit N, Neuhaus K, et al. Sequence and cultivation study of Muribaculaceae reveals novel species, host preference, and functional potential of this yet undescribed family. Microbiome. 2019;7:28.

71. Nagpal R, Wang S, Solberg Woods LC, Seshie O, Chung ST, Shively CA, et al. Comparative Microbiome Signatures and Short-Chain Fatty Acids in Mouse, Rat, Non-human Primate, and Human Feces. Frontiers in Microbiology [Internet]. 2018 [cited 2019 Apr 21];9. Available from: https://www.frontiersin.org/article/10.3389/fmicb.2018.02897/full

72. Ormerod KL, Wood DLA, Lachner N, Gellatly SL, Daly JN, Parsons JD, et al. Genomic characterization of the uncultured Bacteroidales family S24-7 inhabiting the guts of homeothermic animals. Microbiome [Internet]. 2016 [cited 2019 Apr 21];4. Available from: http://microbiomejournal.biomedcentral.com/articles/10.1186/s40168-016-0181-2

73. Smith BJ, Miller RA, Ericsson AC, Harrison DC, Strong R, Schmidt TM. Changes in the gut microbiome and fermentation products concurrent with enhanced longevity in acarbose-treated mice. BMC Microbiol. 2019;19:130.

74. Bruce-Keller AJ, Salbaum JM, Luo M, Blanchard E, Taylor CM, Welsh DA, et al. Obese-type Gut Microbiota Induce Neurobehavioral Changes in the Absence of Obesity. Biological Psychiatry. 2015;77:607–15.

75. Gacias M, Gaspari S, Santos P-MG, Tamburini S, Andrade M, Zhang F, et al. Microbiota-driven transcriptional changes in prefrontal cortex override genetic differences in social behavior. eLife. 2016;5:e13442.

76. Li N, Wang X, Sun C, Wu X, Lu M, Si Y, et al. Change of intestinal microbiota in cerebral ischemic stroke patients. BMC Microbiol. 2019;19:191.

77. Xu R, Tan C, Zhu J, Zeng X, Gao X, Wu Q, et al. Dysbiosis of the intestinal microbiota in neurocritically ill patients and the risk for death. Crit Care. 2019;23:195.

78. Biagi E, Franceschi C, Rampelli S, Severgnini M, Ostan R, Turroni S, et al. Gut Microbiota and Extreme Longevity. Current Biology. 2016;26:1480–5.

79. Liu Y, Luo S, Kou L, Tang C, Huang R, Pei Z, et al. Ischemic stroke damages the intestinal mucosa and induces alteration of the intestinal lymphocytes and CCL19 mRNA in rats. Neuroscience Letters. 2017;658:165–70.

80. Xu X, Zhu Y, Chuai J. Changes in Serum Ghrelin and Small Intestinal Motility in Rats with Ischemic Stroke. The Anatomical Record: Advances in Integrative Anatomy and Evolutionary Biology. 2012;295:307–12.

81. Chen Y, Liang J, Ouyang F, Chen X, Lu T, Jiang Z, et al. Persistence of Gut Microbiota Dysbiosis and Chronic Systemic Inflammation After Cerebral Infarction in Cynomolgus Monkeys. Front Neurol. 2019;10:661.

82. Stanley D, Mason LJ, Mackin KE, Srikhanta YN, Lyras D, Prakash MD, et al. Translocation and dissemination of commensal bacteria in post-stroke infection. Nature Medicine. 2016;22:1277–84.

83. Houlden A, Goldrick M, Brough D, Vizi ES, LénÁrt N, Martinecz B, et al. Brain injury induces specific changes in the caecal microbiota of mice via altered autonomic activity and mucoprotein production. Brain, Behavior, and Immunity. 2016;57:10–20.

84. Xie Q, Pan M, Huang R, Tian X, Tao X, Shah NP, et al. Short communication: Modulation of the small intestinal microbial community composition over short-term or long-term administration with Lactobacillus plantarum ZDY2013. Journal of Dairy Science. 2016;99:6913–21.

85. Sun J, Wang F, Ling Z, Yu X, Chen W, Li H, et al. Clostridium butyricum attenuates cerebral ischemia/reperfusion injury in diabetic mice via modulation of gut microbiota. Brain Res. Netherlands; 2016;1642:180–8.

86. Davari S, Talaei SA, Alaei H, Salami M. Probiotics treatment improves diabetes-induced impairment of synaptic activity and cognitive function: behavioral and electrophysiological proofs for microbiome-gut-brain axis. Neuroscience. United States; 2013;240:287–96.

87. D’Mello C, Ronaghan N, Zaheer R, Dicay M, Le T, MacNaughton WK, et al. Probiotics Improve Inflammation-Associated Sickness Behavior by Altering Communication between the Peripheral Immune System and the Brain. Journal of Neuroscience. 2015;35:10821–30.

88. Kim CH, Park J, Kim M. Gut Microbiota-Derived Short-Chain Fatty Acids, T Cells, and Inflammation. Immune Netw. 2014;14:277.

89. Wang Z, Zhao Y. Gut microbiota derived metabolites in cardiovascular health and disease. Protein Cell. 2018;9:416–31.

90. Yamashiro K, Tanaka R, Urabe T, Ueno Y, Yamashiro Y, Nomoto K, et al. Gut dysbiosis is associated with metabolism and systemic inflammation in patients with ischemic stroke. Smidt H, editor. PLoS ONE. 2017;12:e0171521.

91. Sadler R, Cramer JV, Heindl S, Kostidis S, Betz D, Zuurbier KR, et al. Short-Chain Fatty Acids Improve Poststroke Recovery via Immunological Mechanisms. J Neurosci. 2020;40:1162–73.

92. Chen R, Xu Y, Wu P, Zhou H, Lasanajak Y, Fang Y, et al. Transplantation of fecal microbiota rich in short chain fatty acids and butyric acid treat cerebral ischemic stroke by regulating gut microbiota. Pharmacol Res. 2019;148:104403.

93. Kashyap PC, Chia N, Nelson H, Segal E, Elinav E. Microbiome at the Frontier of Personalized Medicine. Mayo Clinic Proceedings. 2017;92:1855–64.

