## Supplementary figures and images for "Experimental Stroke Induces Chronic Gut Dysbiosis and Neuroinflammation in Male Mice"

### Supplemental Figure 1

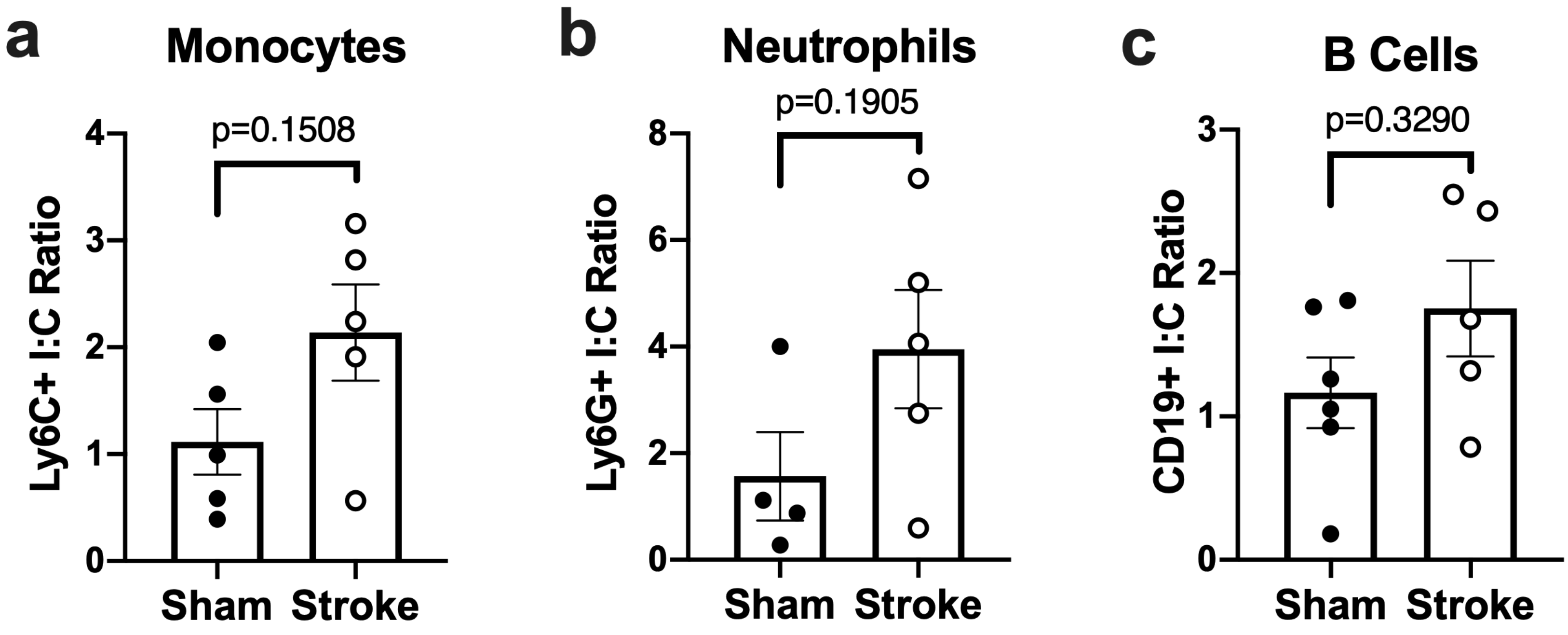

### Supplemental Figure 2

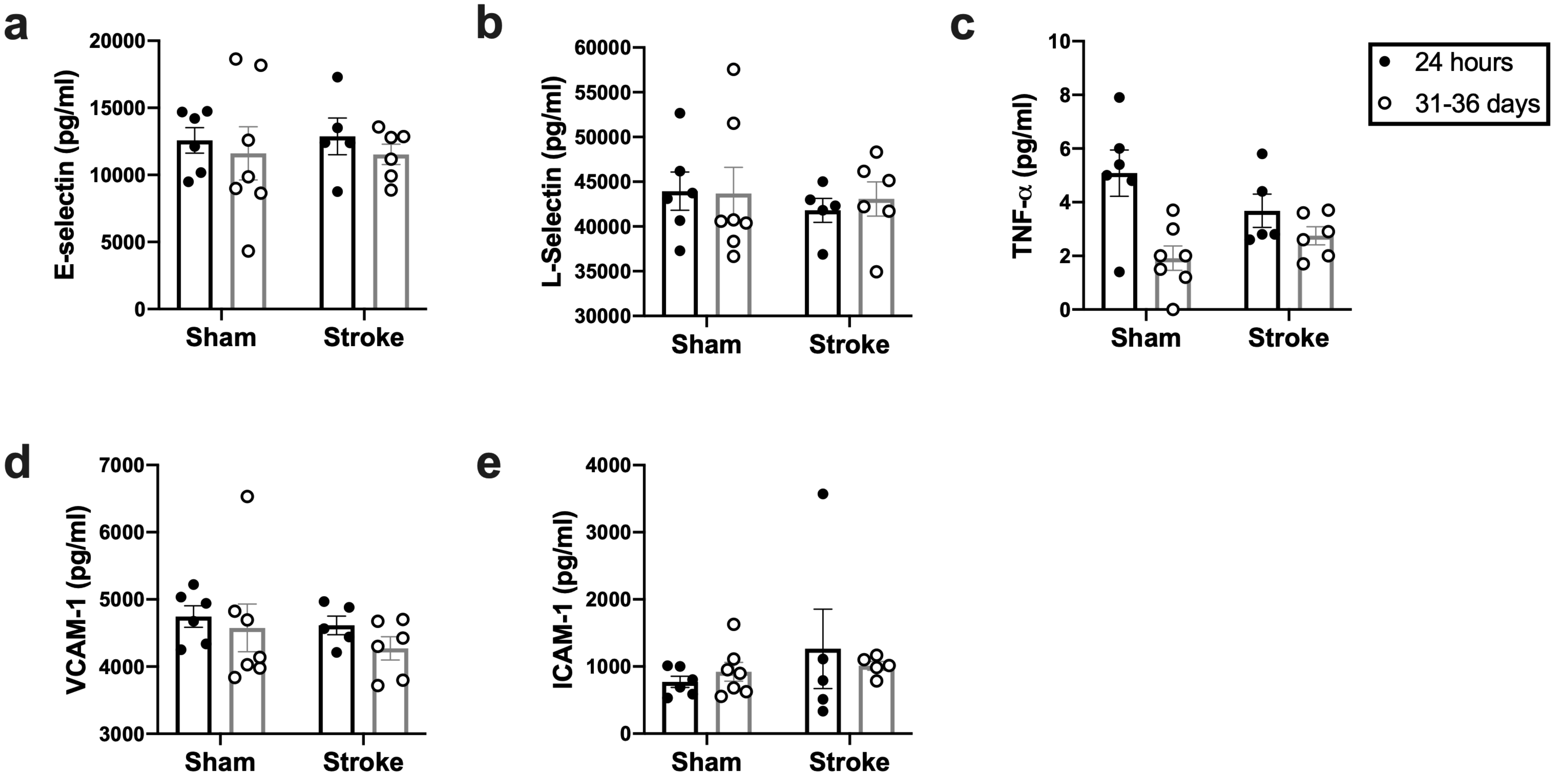
