## Supplemental Tables 1 and 2 for "Experimental Stroke Induces Chronic Gut Dysbiosis and Neuroinflammation in Male Mice"

**Supplemental Table 1: Top reported taxa and their adjusted statistical p-values calculated by LMER.** Bolded taxa and p-values are shown in Figure 8.

|  | **Main Effects** | | **Interaction** |
| --- | --- | --- | --- |
| **Taxa** | **tMCAO** | **Time** | **tMCAO x Time** |
| **Phylum** | | | |
| **Bacteroidetes** | 0.92 | 0.88 | **0.07** |
| **Firmicutes** | 0.92 | 0.88 | **0.03** |
| Acidobacteria | 0.92 | 0.09 | 0.41 |
| Actinobacteria | 0.92 | 0.88 | 0.41 |
| Proteobacteria | 0.36 | 0.30 | 0.29 |
| **Class** | | | |
| **Bacteroidia** | 0.88 | 0.88 | **0.07** |
| **Clostridia** | 0.78 | 0.88 | **0.05** |
| Bacilli | 0.87 | 0.88 | 0.70 |
| Gammaproteobacteria | 0.24 | 0.88 | 0.32 |
| Actinobacteria | 0.87 | 0.88 | 0.49 |
| **Order** | | | |
| **Bacteroidales** | 0.88 | 0.88 | **0.07** |
| **Clostridiales** | 0.78 | 0.88 | **0.05** |
| Lactobacillales | 0.78 | 0.88 | 0.76 |
| Enterobacteriales | 0.23 | 0.88 | 0.34 |
| Bifidobacteriales | 0.78 | 0.88 | 0.58 |
| **Family** | | | |
| **S24-7*/Muribaculaceae*** | 0.76 | 0.93 | **0.03** |
| Lachnospiraceae | 0.76 | 0.93 | 0.06 |
| ***Ruminococcaceae*** | 0.76 | 0.93 | **0.03** |
| *Lactobacillaceae* | 0.76 | 0.93 | 0.76 |
| **Genus** | | | |
| ***Oscillospira*** | 0.66 | 0.82 | **0.03** |
| *Lactobacillus* | 0.79 | 0.82 | 0.94 |
| *Coprococcus* | 0.99 | 0.82 | 0.94 |

**Supplemental Table 2: Top reported taxa and their adjusted statistical p-values calculated by ZIBR.** P-values <0.05 are in bold.

|  | **Main Effects** | | **Interaction** |
| --- | --- | --- | --- |
| **Taxa** | **tMCAO** | **Time** | **tMCAO x Time** |
| **Phylum** | | | |
| Bacteroidetes | 0.99 | 0.985 | 0.057 |
| Firmicutes | 0.74 | 0.984 | **0.015** |
| Proteobacteria | 0.76 | 0.115 | 0.793 |
| Actinobacteria | 0.89 | 0.840 | 0.786 |
| Acidobacteria | 0.55 | **0.006** | 0.612 |
| **Class** | | | |
| Bacteroidia | 0.99 | 0.984 | 0.058 |
| Clostridia | 0.53 | 0.993 | **0.026** |
| Bacilli | 0.87 | 0.424 | 0.530 |
| Gammaproteobacteria | 0.41 | 0.132 | 0.322 |
| Actinobacteria | 0.89 | 0.634 | 0.724 |
| Epsilonproteobacteria | 0.27 | **0.028** | 0.133 |
| Betaproteobacteria | 0.42 | **0.000003** | **0.0003** |
| **Order** | | | |
| Bacteroidales | 0.99 | 0.98 | 0.058 |
| Clostridiales | 0.53 | 0.99 | **0.026** |
| Lactobacillales | 0.72 | 0.34 | 0.50 |
| Enterobacteriales | 0.31 | 0.11 | 0.22 |
| Bifidobacteriales | 0.57 | 0.67 | 0.69 |
| Campylobacterales | 0.27 | **0.028** | 0.13 |
| Burkholderiales | 0.37 | **0.000003** | **0.00003** |
| **Family** | | | |
| S24-7*/Muribaculaceae* | 0.92 | 0.99 | **0.009** |
| *Lachnospiraceae* | 0.89 | 0.92 | 0.13 |
| *Lactobacillaceae* | 0.74 | 0.35 | 0.52 |
| *Ruminococcaceae* | 0.86 | 0.81 | 0.13 |
| *Turicibacteraceae* | 0.39 | 0.12 | 0.086 |
| **Genus** | | | |
| *Lactobacillus* | 0.74 | 0.35 | 0.52 |
| *Oscillospira* | 0.75 | 0.87 | 0.075 |
| *Coprococcus* | 0.99 | 0.83 | 1.0 |
| *Enterobacter* | **0.019** | 0.51 | **0.007** |
| *Bifidobacterium* | 0.57 | 0.67 | 0.69 |
